# Modeling Thick Filament Activation Suggests a Molecular Basis for Force Depression

**DOI:** 10.1101/2023.09.27.559764

**Authors:** Shuyue Liu, Chris Marang, Mike Woodward, Venus Joumaa, Tim Leonard, Brent Scott, Edward Debold, Walter Herzog, Sam Walcott

## Abstract

Multiscale models aiming to connect muscle’s molecular and cellular function have been difficult to develop, in part, due to a lack of self-consistent multiscale data. To address this gap, we measured the force response from single skinned rabbit psoas muscle fibers to ramp shortenings and step stretches performed on the plateau region of the force-length relationship. We isolated myosin from the same muscles and, under similar conditions, performed single molecule and ensemble measurements of myosin’s ATP-dependent interaction with actin using laser trapping and in vitro motility assays. We fit the fiber data by developing a partial differential equation model that includes thick filament activation, whereby an increase in force on the thick filament pulls myosin out of an inhibited state. The model also includes a series elastic element and a parallel elastic element. This parallel elastic element models a titin-actin interaction proposed to account for the increase in isometric force following stretch (residual force enhancement). By optimizing the model fit to a subset of our fiber measurements, we specified seven unknown parameters. The model then successfully predicted the remainder of our fiber measurements and also our molecular measurements from the laser trap and in vitro motility. The success of the model suggests that our multiscale data are self-consistent and can serve as a testbed for other multiscale models. Moreover, the model captures the decrease in isometric force observed in our muscle fibers after active shortening (force depression), suggesting a molecular mechanism for force depression, whereby a parallel elastic element combines with thick filament activation to decrease the number of cycling cross-bridges.

**SIGNIFICANCE:** Connecting the molecular and cellular scales of muscle contraction would assist in, e.g., the treatment of genetic muscle diseases, the development of heart drugs, and the design of prostheses. The history dependence of muscle contraction, having no clear molecular basis, has remained an obstacle in making this connection for the seventy years since its discovery. We measured the force- and motion-generating capacity of rabbit psoas muscle from the scale of single molecules to single cells. We developed a mathematical model that, when fit to some of the cellular measurements, predicted the remaining cellular measurements and also the molecular measurements. The model’s ability to capture muscle’s history dependence suggests a unified description of muscle contraction from the molecular to cellular scale.

## INTRODUCTION

With the remarkable advances of molecular biology and biophysics over the past 40-50 years has come the increasing need to understand how molecular-scale events give rise to processes observed at the cellular and tissue level. For example, ever since the groundbreaking measurements of a single, isolated myosin molecule interacting with an actin filament (1–4) researchers have worked to relate molecular properties, including those that give rise to devastating diseases, to muscle function *in vivo* (5–11). Mathematical and computational modeling has been critical to this process, particularly in providing insight into emergent properties, the often unexpected behaviors that arise when individual entities work collectively (12–15). Numerous models have been developed that fit measurements at one scale and make plausible predictions of measurements at another (14, 16–20); however, as far as we are aware, no single model exists that is capable of *quantitatively* fitting muscle measurements from the molecular to cellular scale.

The inability of models to span the molecular to cellular scales is not due to a lack of clarity at the molecular level. On the contrary, though some details of myosin’s cross-bridge cycle remain unclear (e.g. the sequencing of the power-stroke with phosphate release (21–24) and the reversibility of the power-stroke (25)), most of the biochemical states and their sequence has been known for decades (26). A more recent advance in understanding myosin’s interaction with actin has been the direct measurement of myosin’s force dependent kinetics, which had been predicted long before it was measured (27, 28). The force dependence of myosin means that, in addition to the biochemical states of myosin’s interaction with actin, mathematical models must also define how force influences the rate constants governing transitions between the states. Lacking direct measurements, early models had the freedom to choose functions defining this force dependence based on convenience or computational efficiency (28–32). However, many reaction rates have now been directly or indirectly measured in the presence of an applied force (33–37) constraining the functional form of the force-dependence, reducing the number of model parameters, and making it possible to perform parameter estimation by optimizing model fits to data. It is therefore, we believe, an opportune time to develop a molecular-to-cellular muscle model.

A multiscale muscle model does not yet exist, in part, due to the fact that (as far as we are aware) no self-consistent multiscale data set exists. That is, we are unaware of measurements of actomyosin interaction, spanning the molecular to cellular scale and collected under the same conditions with myosin from the same tissue. This lack of self-consistency introduces uncertainties in model parameters (e.g., the rates of biochemical state transitions) caused by using different proteins and/or different experimental conditions. This kind of uncertainty makes it difficult to construct and test a multiscale model capable of explaining the emergent and often complex properties of muscle, such as its history dependence. In particular, any potential model must account for both the decrease in isometric force following muscle shortening (force depression) and the increase in isometric force following muscle lengthening (force enhancement), neither of which has a widely-accepted molecular basis (38, 39). These properties of muscle present a particularly strong challenge, and test, for any multiscale model. Therefore, the purpose of this study was two-fold: first, to collect a multiscale data set to serve as a test-bed for any putative multiscale muscle model; second, to present a minimal model that is able to explain the data. In doing so, we show not only that the data are described by the same model with the same parameters and are therefore self-consistent, but we also propose a plausible model for muscle’s history dependence.

## RESULTS AND DISCUSSION

We performed measurements on single skinned muscle fibers and on purified myosin *in vitro* isolated from the same rabbit psoas muscle for all experiments. As much as possible, we matched buffers and experimental conditions. Our experimental measurements were selected to span a wide range of length-time histories (at the fiber scale) and ATP concentrations and ensemble sizes (at the molecular scale). Specifically, each skinned fiber was activated at a length on the plateau region of the force-length relationship, and then subjected to a force-velocity protocol (5% ramp shortening over a range of times, from 0.01s to 30s) and a quick stretch protocol (0.5% and 0.75% stretches). Experiments were performed with purified myosin using single molecules and mini ensembles in the laser trap, and in the in vitro motility assays. We purposely chose experiments routinely performed by our labs to minimize experimental error, the novelty lies in the self-consistency of the measurements from the single molecule to cellular scales (see Methods for details).

### Muscle fiber experiments

With one notable exception, our measurements of muscle fiber force for ramp shortening and step lengthening were consistent with previous observations (Fig. 1). Force initially dropped steeply during the shortening ramp and then more slowly, without reaching a steady-state (40). During the isometric hold after stretch, force slowly recovered to a lower value than the pre-shortening isometric force (Fig. 1A). Since shortening is done over the plateau region of the force-length relationship, we assume fiber length has no effect on isometric force and this observed decrease following stretch is force depression (41). Force peaks during the step increase in length and then slowly decreases during the subsequent isometric hold. We would expect to observe residual force enhancement (41), i.e. that the force during this post-stretch isometric phase would approach a higher value than the pre-stretch isometric force; however, we performed a shortening step 0.1s after the lengthening step, which eliminated force enhancement, and force returned to the pre-stretch isometric value (Fig. 1B, see Supplementary Material, SM). To quantify force depression in the ramp shortening experiments, we determined force at the end of the ramp shortening, and 0.1, 1 and 2 seconds after (Fig. 2). Strikingly, we observe that any dependence on the duration of the shortening ramp (or, equivalently, shortening velocity) is eliminated after 2s (no significant difference, *p* = 0.16, one-way ANOVA), with force only 50.8±0.05% (mean±SD) of the pre-stretch value. This finding contrasts with previous measurements of force depression that have observed a decrease in force depression with increasing shortening velocity (42–46).

**Figure 1.**
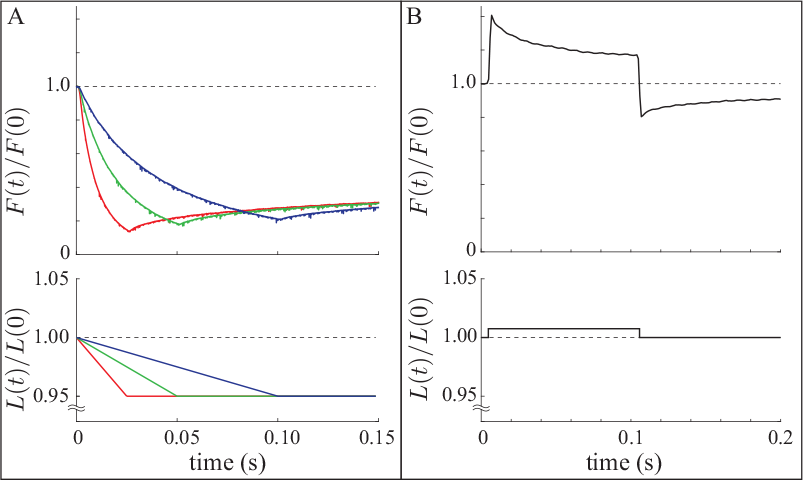
Example measurements from a single skinned fiber. A. In the ramp shortening experiments, fibers were shortened by 5% of their length at a constant rate with the shortening applied over variable times (bottom, 25ms, 50ms and 100ms shown). The force response (top, shown relative to isometric) was recorded for *n* = 8 fibers. B. In the step stretch experiments, fibers were rapidly lengthened, held for 0.1s and then shortened back to their original length (bottom, 0.75% shown). The force response (top, shown relative to isometric) was recorded for *n* = 8 fibers.

**Figure 2.**
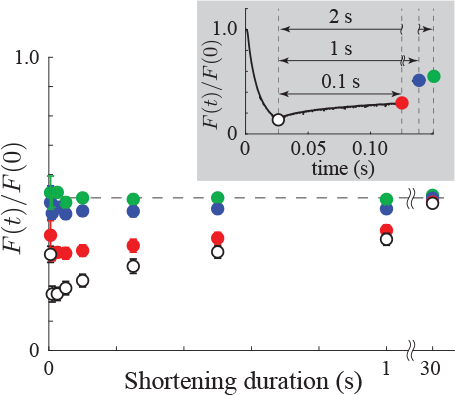
Force after shortening is velocity independent. Force after the end of the ramp shortening (mean±SEM, *n* = 8) shows a clear dependence on shortening duration for < 1s, but that dependence disappears over longer times (green dots). Since each fiber is shortened the same distance, the different shortening durations correspond to different shortening velocities. Note, *t* = 30s has *n* = 3, *t* = 0.01s has *n* = 2.

### Molecular experiments

Our single molecule measurements of myosin’s step size (*d*) performed with the three-bead laser trap assay (Fig. 3) show no dependence on ATP concentration, while measurements of attachment duration (*t*_*on*_) show the expected approximate hyperbolic dependence (1). Interestingly, we find a decreased step size, *d* = 5.27±0.34nm, (mean±SEM of all events *n* = 1198) and an increased apparent ATP binding rate, *k*_*T*_ = 4.0±1.0*μ*M^−1^s^−1^, (mean ± SEM, from applying the equation *t*_*on*_ = 1/*k*_*T*_ [ATP] to average event lifetime at each [ATP]) compared to the values we have previously measured with chicken skeletal myosin, *d*≈10nm and *k*_*T*_≈2*μ*M^−1^s^−1^ (14, 47). Due to this discrepancy, we used our other molecular measurements to estimate step size as independent tests of this finding.

**Figure 3.**
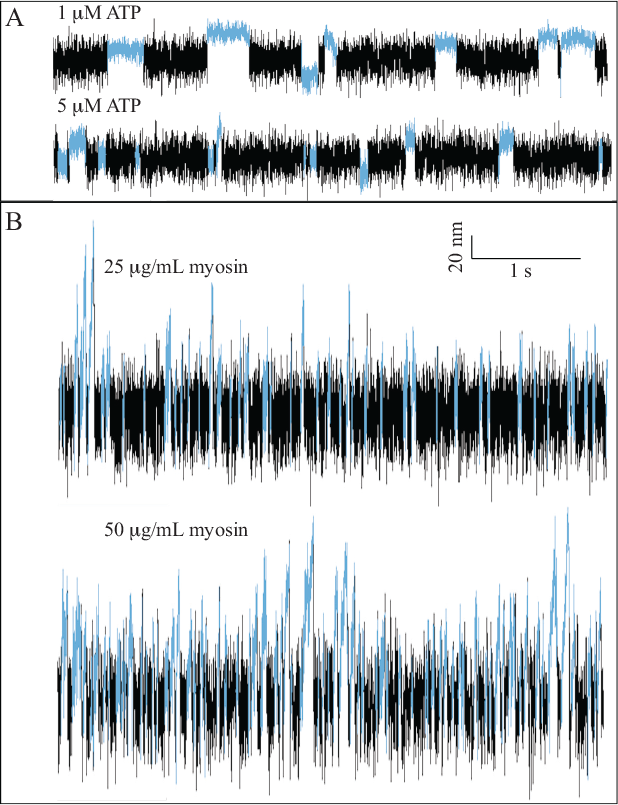
Example displacements of the trapped bead in the laser trap assay. A. Bead position under single molecule conditions at different ATP concentrations. Sections of the trace identified as binding events are colored blue. B. Bead position under mini ensemble conditions at different myosin concentrations and 100 *μ*M ATP. Sections of the trace identified as binding events are colored in blue. Scale bars are the same for all plots.

Our mini-ensemble measurements of maximum displacement (*d*_*max*_) and event duration (*t*_*event*_), performed at an ATP concentration of 100*μ*M show an increase in both *d*_*max*_ and *t*_*event*_ with increasing myosin (see Table 1, Fig. 3). From these measurements, we can estimate myosin’s step size, *d*, by dividing *d*_*max*_ by *t*_*event*_ to get an effective speed (*v* = 1.90*μ*m s and *v* = 2.01*μ*m s for 25 and 50*μ*g/mL, respectively) and multiplying by the expected attachment time (1/*k*_*T*_ [ATP] = 2.5ms, given *k*_*T*_ = 4.0*μ*M^−1^s^−1^ and [ATP] = 100*μ*M). This calculation estimates a unitary step size of 4.8±1.2nm (25*μ*g/mL, error propagated using SEM) and 5.0±1.3nm (50*μ*g/mL, error propagated using SEM), consistent with the reduced step size we directly observed at the single molecule level.

**Table 1:** Mini-ensemble measurements.

| Myosin concentration ($\mu\text{g/mL}$) | $d_{\text{max}}$ (nm) | $t_{\text{event}}$ (ms) |
| --- | --- | --- |
| 25 ($n = 617$) | $38.22 \pm 0.70$ | $20.08 \pm 0.45$ |
| 50 ($n = 813$) | $47.12 \pm 0.76$ | $23.41 \pm 0.73$ |

Numbers listed are mean plus/minus SEM.

Finally, our *in vitro* motility measurements show actin or regulated thin filament (RTF) speeds of 4-5 *μ*m s at saturating ATP. RTF speed was a bit higher than actin speed at high ATP, but the difference was small (a Michaelis-Menten fit gives *V*_*max*_ = 4.21*μ*m/s and *K*_*m*_ = 81.6*μ*M for actin and *V*_*max*_ = 5.03*μ*m/s and *K*_*m*_ = 131*μ*M for RTFs). From these measurements, we can estimate myosin’s step size, *d* by fitting a straight line constrained to pass through the origin to data at low [ATP] ([ATP] ≤100*μ*M), which gives a slope of 0.0297 (actin) and 0.0258*μ*m/s*μ*M (RTF). Dividing these by the ATP binding rate (*k*_*T*_ = 4.0*μ*M^−1^s^−1^) gives an estimated step size of *d* = 7.4±2.1 for actin and *d* = 6.5± 2.0nm for RTFs (error propagated using SEM). As with the mini-ensemble measurements, the motility measurements are consistent with the step size we observed in the single molecule laser trap assay.

### Mathematical model

Our observation that force depression (the force 2s after the end of the ramp stretch) is independent of shortening rate (Fig. 2, green dots) simplifies the modeling considerably since, if force depression arises in whole or in part from a decrease in cycling cross-bridges (43, 46), then cross-bridges only need to sense sarcomere length as opposed to both length and shortening velocity (or, e.g., work done by the fiber (42)). Given that titin has already been proposed to bind to actin during muscle lengthening (48), titin seems a likely means to sense sarcomere length during shortening and a force-dependent transition into an inhibited state (49–51) a likely mechanism to amplify the drop in force during shortening, and explain why the number of cycling cross-bridges decreases during force depression.

To test this idea, we implemented it into a mathematical model. The model contains three components: 1) a series spring to model all elastic structures in series with actomyosin (e.g., the z-disc, thick and thin filaments, non-functional sarcomeres at the end of the muscle fiber, elasticity in the experimental apparatus, etc); 2) a parallel spring to model elastic structures in parallel with actomyosin (particularly the hypothesized interaction of titin with actin); and 3) a contractile element that accounts for the molecular interactions of myosin with actin (Fig. 4). This arrangement is similar to that in Hill-models (52), that are often used to capture the behavior of whole muscle. However, in our model we are explicitly modeling molecular structures.

**Figure 4.**
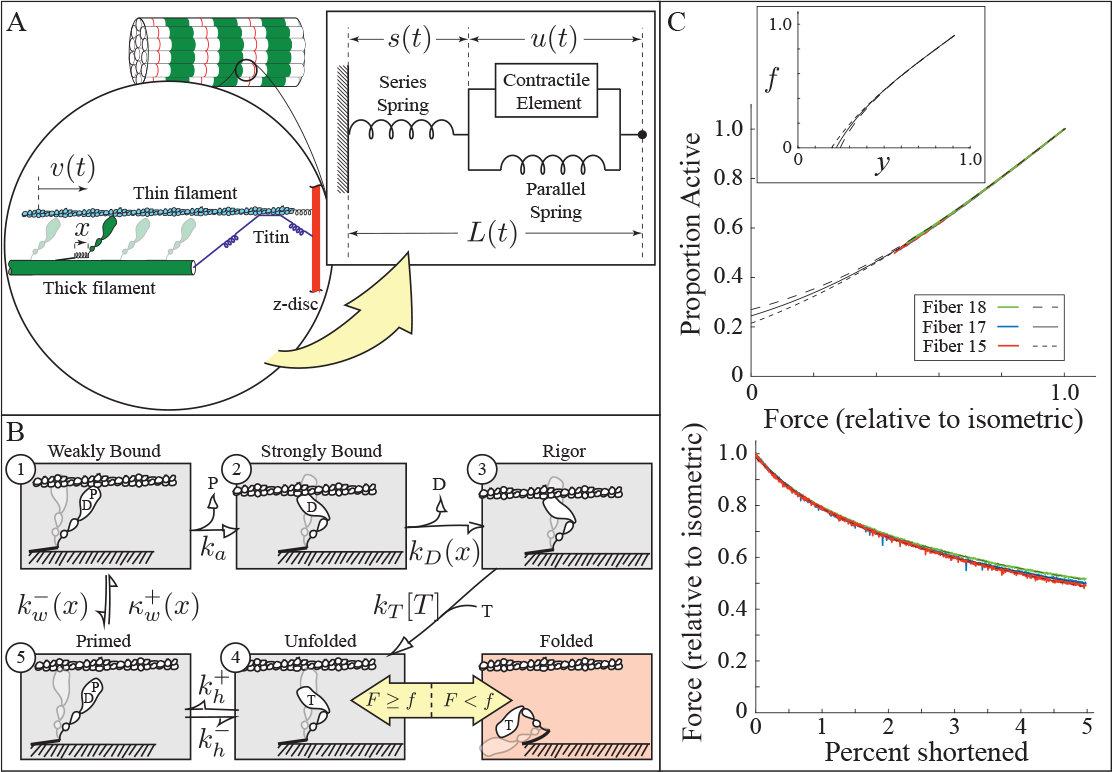
Model schematics. A. A half sarcomere is modeled as elastic elements in series and parallel with the contractile element, wherein the force of myosin interacting with actin is generated. The parallel spring models an interaction between titin and actin. B. Five state kinetic model for active myosin (gray), and a folded, inactive state (red). C. Force measurements for slow (30s) shortening ramps (bottom) provide estimates of active, unfolded myosin as a function of thick filament force (top); the inverse of this function is the folding threshold, *f* (*y*), inset. Abbreviations: D is ADP, P is inorganic phosphate, T is ATP.

For the contractile element, we use a five-state model to capture myosin’s interaction with actin (Fig. 4A). This model is defined by the following integro-partial differential equations (iPDEs) (53):

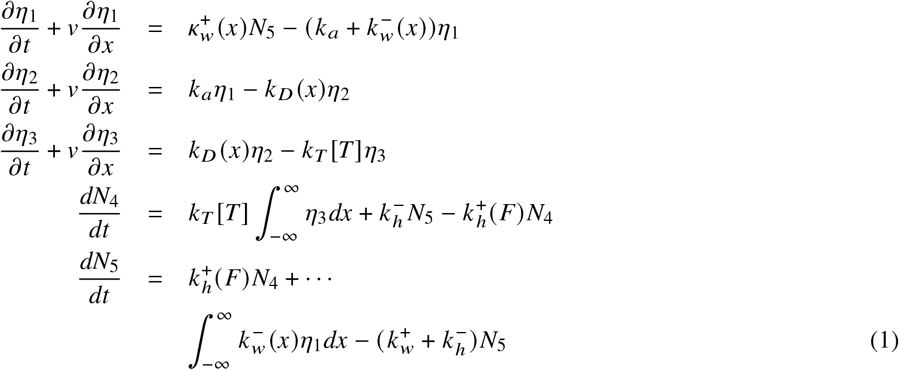

where *η*_*i*_ (*x, t*) is a probability density, *N*_*i*_ (*t*) is a probability (such that 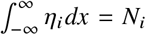), *κ* _*j*_ (*x*) is a rate density, and *k* _*j*_ is a rate (such that 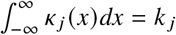). Note that 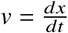 is the relative velocity of actin past the thick filament, *x* is the extension of myosin, and *t* is time. There is an additional constraint of conservation of probability, *N*_1_ + *N*_2_ + *N*_3_ + *N*_4_ + *N*_5_ = 1. The contractile element generates force as 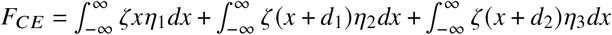, where ζ is the stiffness of myosin, *d*_1_ is the powerstroke that occurs during the transition from state 1 to 2, and *d*_2_ is *d*_1_ plus the additional step that occurs during the transition from state 2 to 3.

The input to the model is total muscle length as a function of time, *L* (*t*), which is made up of the extension of the series spring (*s*(*t*)) and the accumulated lengthening of the contractile element 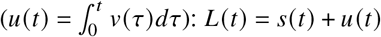. Then, since the contractile element and parallel spring are in series with the series spring, their forces are equal:

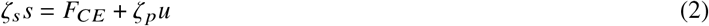

where ζ_*s*_ is the stiffness of the series spring and ζ _*p*_ is the stiffness of the parallel spring.

Two rate constants depend on molecular extension, 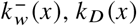, *k*_*D*_ (*x*). The former arises because weakly-bound myosin can be ripped off, and has the form 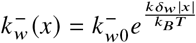, where *k*_*w*0_ is the reaction rate at zero extension and δ_*w*_ determines the force sensitivity of this reaction (35). The latter arises because ADP release is force dependent, and has the form 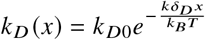 (33–37). The weak binding rate density, 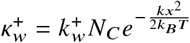, where *N*_*C*_ is the normalization constant such that 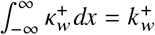.

To model thick filament activation (49–51), we make the rate constant 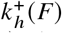 a function of the force on the thick filament, *F* = *F*_*CE*_ + ζ _*p*_*u*. If thick filament force drops below a threshold, *F f y*, then a given myosin molecule will fold up and not hydrolyze ATP once it enters the unbound, pre-hydrolysis state (state 4 in the model, Fig. 4B). If force is then increased, *F* ≥ *f* (*y*), then that myosin molecule extends and can hydrolyze ATP, bind actin, and continue through its cycle.

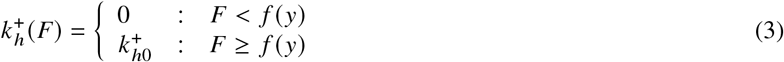

We assume that each myosin molecule may have a different threshold, which we implement with the function *f* (*y*). The variable 0≤ *y*≤ 1 gives the proportion of myosin in the ensemble with a force threshold less than or equal to *f* (*y*) – that is, *f*(0.3) = 0.1*F*_*iso*_, would indicate that 30% of the myosin molecules in a given ensemble have an activation threshold less than or equal to one tenth of the isometric force, *F*_*iso*_. By implementing thick filament activation as described, as opposed to a force-dependent equilibrium constant, the model produces two distinct ATP turnover rates since there is no interchange between the folded and unfolded state at a given force (54, 55).

The model has the following parameters, *k*_*T*_, *d*_1_, *d*_2_, 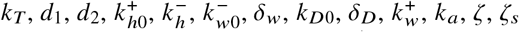, δ_*w*_, *k*_*D*0_, δ_*D*_, 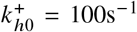, *k* _*a*_, ζ, ζ_*s*_, and ζ _*p*_. Additionally, the model requires the specification of the function *f*(*y*). We assume 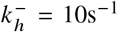 (56), (56), *d*_2_−*d*_1_ = 0 (for simplicity), and ζ = 1pN/nm (57, 58). From measurements of force enhancement as a function of stretch amplitude (59), we estimate a net stiffness of 0.1% isometric force/nm, which allows us to specify ζ _*p*_ given ζ_*s*_. From our single molecule measurements, we estimate *d*_1_ = 6nm and *k*_*T*_ = 4*μ*M^−1^s^−1^. Finally, we can estimate *f*(*y*) by assuming that our slowest ramp shortening experiments, where shortening occurred over 30s, are in quasi steady-state (Fig. 4C, see Methods). The model therefore has seven free parameters (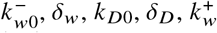, δ_*w*_, *k*_*D*0_, δ_*D*_, 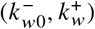, *k* _*a*_, and ζ_*s*_) that we varied to optimize the fit of the model to our skinned fiber measurements (see Methods).

### Model fits

We fit the model to 0.15s of force measurements for two ramp shortenings (0.025s and 0.1s) and 0.2s of force measurements for one step stretch (0.75% length increase). We performed these fits for three different skinned fibers (Fig. 5, red curves, SM for all fits), and the resulting best-fit parameters are given in Table 2 (see SM for a sensitivity analysis).

**Table 2:** Best fit parameters of model fits to three fiber measurements.

| Fiber number | 15 | 17 | 18 |
| --- | --- | --- | --- |
| $k_{D0}$ ($s^{-1}$) | 3130 | 7000 | 7300 |
| $\delta_D$ (nm) | 1.22 | 1.51 | 1.61 |
| $k_{w0}^-$ ($s^{-1}$) | 13100 | 7460 | 12600 |
| $\delta_w$ (nm) | 0.135 | 1.31 | 3.54 |
| $k_w^+$ ($s^{-1}$) | 2760 | 3590 | 12600 |
| $k_a$ ($s^{-1}$) | 455 | 297 | 296 |
| $\zeta_s$ (pN/nm/myosin) | 0.0265 | 0.0159 | 0.0208 |

**Figure 5.**
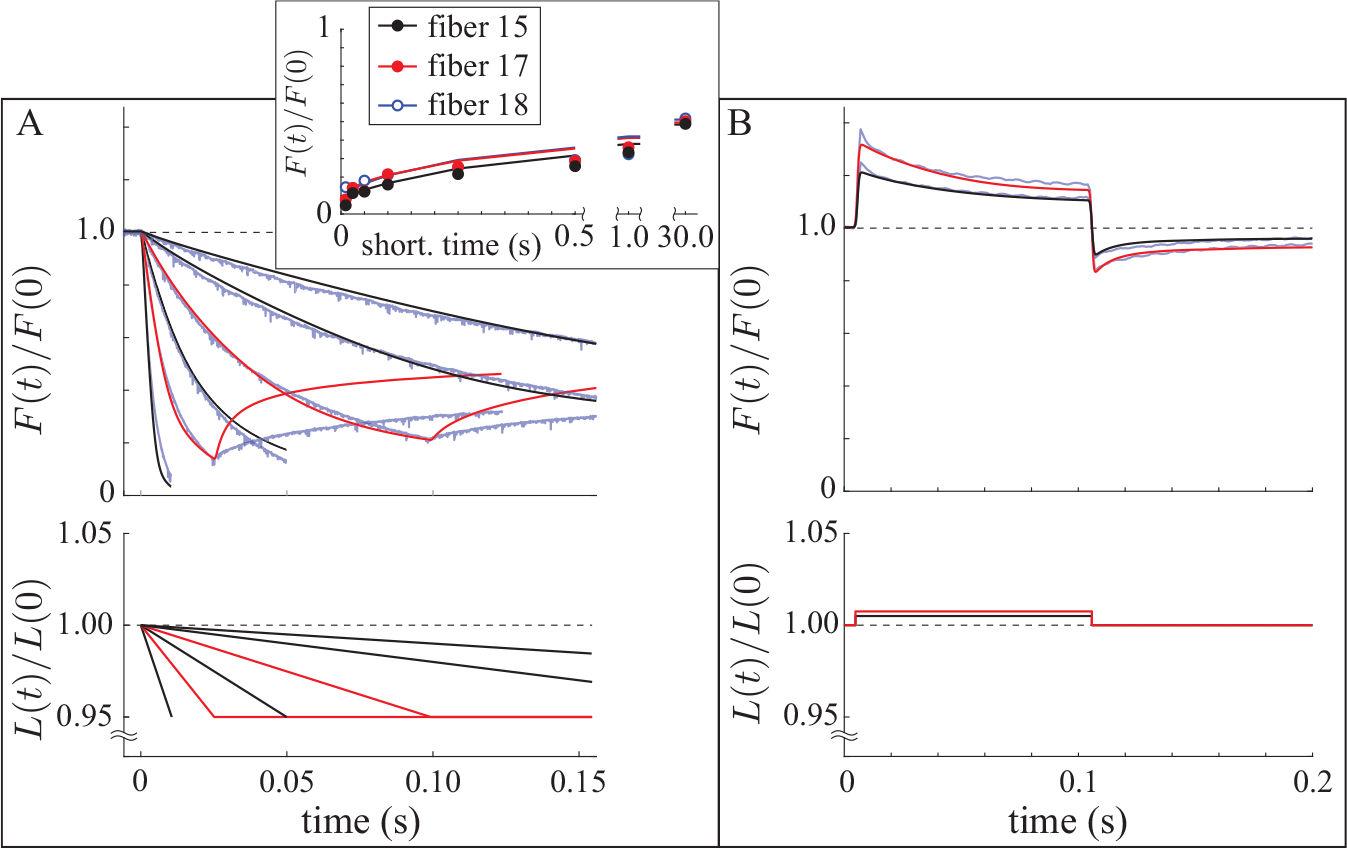
Model fits and predictions. In all plots, red curves are fits, black curves are model predictions, and blue curves are measurements. A. Ramp shortening. The model reasonably captures the force response during shortening, both for the fitted measurements and for the predicted measurements. The model always over-predicts the force rise after shortening, which is shown only for the fitted measurements for visualization purposes. Inset shows the force at the end of the ramp as a function of shortening duration for all fibers. B. Step lengthening/shortening. The model reasonably captures the force response during and after lengthening and shortening. Measurements from fiber 17.

Broadly, we find that these parameters are consistent with, though the ADP release (*k*_*D*0_) and weak binding rates 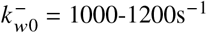 are at the upper end of previous estimates. In particular *k*_*D*0_ has been estimated to be in the range of 100-500s^−1^ (14, 35, 60), though faster estimates exist (e.g., ≥2000s^−1^, (61, 62)). In our experiments, a faster ADP release rate is necessitated, in part, by a smaller myosin step size, since ADP release rate would need to be at least velocity divided by step size, 6*μ*m/s/6nm = 1000s^−1^. Additionally, drag on the thin filament from weakly bound myosin increases the ADP release rate required for a given velocity, since a resistive force both decreases myosin’s step size (63) and slows myosin’s ADP release. Consequently, the slower estimates of ADP release rate are not consistent with our experimental observations, so our model’s estimates are necessarily faster.

The model’s predicted force dependence of ADP release, δ_*D*_, is consistent with measurements in the ultra-fast laser trap of δ_*D*_ = 1.4-1.6nm (35). Indirect measurements cannot generally determine δ_*D*_ directly, but can determine the non-dimensional parameter *E* = ζ δ_*D*_ *d*/*k*_*B*_*T*, where ζ is myosin’s stiffness, *d* is the powerstroke size, and *k* _*B*_*T* is Boltzmann’s constant times temperature (14, 64). From the model fits, we predict *E* = 1.77-2.33 (*E* = 1.77, 2.19, 2.33 for fibers 15, 17, 18, respectively), a bit higher than was predicted from fits of a model to *in vitro* measurements of chicken pectoralis myosin, *E* = 1.35 (14). The values are more consistent with estimates from Fenn’s heat of shortening measurements with frog muscle, *E* = 1.88 (27, 53).

Weak binding is a challenge to measure directly. In one of the few measurements of which we are aware, Capitanio et al. (35) measure 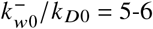 and δ_*w*_ = 1.3-1.4nm. While this weak binding rate is much less than predicted by the model, so too was the ADP release rate from the same experimental data (*k*_*D*0_ = 200s^−1^). Interestingly, the relative values are more similar, 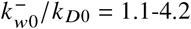 (35) compared to 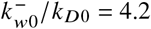 (fiber neq], fiber 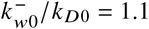, fiber 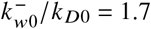), raising the possibility that myosin is overall faster in our experimental conditions.

The weak to strong binding transition, *k*_*a*_, is difficult to measure directly, but is thought to limit ATP hydrolysis. We can estimate the solution ATP hydrolysis rate by calculating the occupancy of each state under steady-state, zero force conditions (see Methods). We predict an ATPase rate, 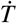, between 40.9 and 57.6s^−1^ (fiber 15 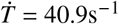, fiber 17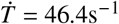, fiber 18 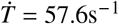), which is consistent with previous measurements of solution ATPase of 50-75s^−1^ (65, 66).

The series (ζ_*s*_) and parallel (ζ _*p*_) stiffnesses in the model are defined per myosin molecule. If we estimate 150 myosin molecules in a half sarcomere, we would estimate ζ_*s*_ = 4.0 pN/nm (fiber 15), ζ_*s*_ = 2.4 pN/nm (17), and ζ_*s*_ = 3.1 pN/nm (18). This stiffness combines all of the elastic structures in series with the skinned fiber preparation, so it is difficult to compare it directly to measured stiffnesses. However, parallel stiffness, which can be calculated from this value (see Methods) is ζ _*p*_ = 0.054 pN/nm (fiber 15), ζ _*p*_ = 0.048 pN/nm (fiber 17), and ζ _*p*_ = 0.055 pN/nm (fiber 18), roughly two orders of magnitude less than ζ_*s*_. This result means that the overall stiffness of the system is approximately the parallel stiffness and, if the parallel spring arises from titin binding to actin (Fig. 4), will reflect titin’s stiffness.

### Model predictions

From these best fits, we then predicted the force response to ramp shortenings of duration ≤ 1s (0.01s, 0.05s, 0.25s, 0.5s 1s) and the force response to a step increase of 0.5%. With the exception of the force response immediately after the shortening ramps, which the model consistently over-predicts, the agreement is good (Fig. 5, see SM for all model fits to/predictions of fiber measurements and an analysis of the model’s predictive ability).

We next used each best-fit parameter set from the skinned fiber measurements to predict our molecular measurements. In these experiments, we expect that the myosin molecules cannot form the self-inhibited folded state (51, 67). Moreover, there is no series or parallel elastic element. Finally, in the laser trap, because the bead-actin-bead “dumbbell” assembly fluctuates in the z direction (i.e., perpendicular to the surface of the flow cell), the attachment of the first myosin molecule is slowed by roughly an order of magnitude but, once that first myosin binds, subsequent myosin bind at the same rate as in a fiber (14, 19). With these assumptions, we used a Monte-Carlo simulation to predict the results of the single molecule and mini-ensemble experiments in the laser trap, and the results of *in vitro* motility (see Methods).

The single molecule measurements reasonably agree with our simulations (Fig. 6). While this result is, perhaps, unsurprising since we used our single molecule measurements to specify the power stroke size (*d*_1_ = 6nm) and ATP binding rate (*k*_*T*_ = 4*μ*M^−1^s^−1^) in the model, we used our model to simulate laser trap data, which we then analyzed with the same programs as our experimental data. In this way, if the weak-binding interactions lasted long enough to be detected, we would observe deviations from the theoretical predictions of *d*_1_ and *t*_*on*_ that become more pronounced at higher ATP where the strong-binding lifetime is decreased. Since we do not observe this, we conclude that weak binding in our model is not detectable in our single molecule laser trap.

**Figure 6.**
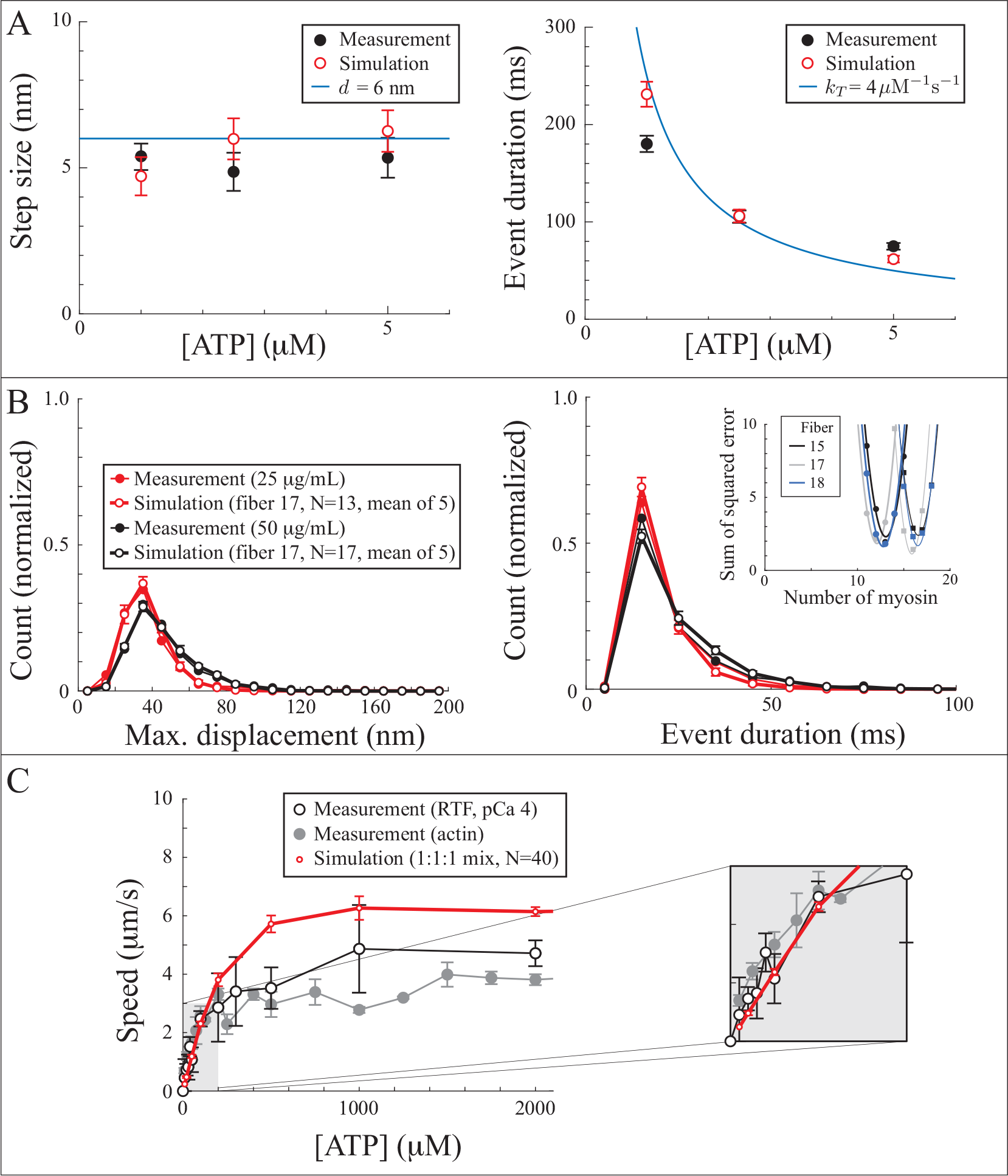
Model predictions agree with molecular measurements. A. Single molecule measurements of step size (*d*) and event duration (*t*_*on*_) agree with model predictions. The model was used to simulate laser trap data (bead position vs. time) and analyzed with the same programs as our measurements. Error bars show SEM. B. Mini ensemble measurements of maximum displacement (*d*_*max*_) and event duration (*t*_*event*_) agree with model predictions. The model was used to simulate laser trap data (bead position vs. time) and analyzed with the same algorithms as our measurements. Symbols show the mean and error bars the standard deviation of five simulations. To determine the number of myosin molecules in the mini ensemble, we calculated the mean squared difference between the simulated and measured cumulative probability distributions for a range of different ensemble sizes (inset, mean of five simulations for myosin density of 25*μ*g mL are circles, myosin density of 50*μ*g mL are squares, lines are quadratic fits). C. In vitro motility measurements with actin and regulated thin filaments (RTFs) reasonably agree with simulations of an ensemble of 40 myosin molecules each with parameters selected with equal probability from one of the three parameter sets in Table 2. Inset shows measurements at low ATP, where detachment is limited by ATP binding.

In the mini ensemble measurements, we do not exactly know the number of myosin molecules present. We therefore performed a series of simulations with different ensemble sizes (see Methods). In these simulations, we again simulate the laser trap data and then analyze it with the same algorithms as our experimental data. Each of the three best-fit parameter sets reasonably agreed with our measurements and predict *n* = 12-13 and *n* = 16-17 for myosin concentrations of 25 and 50*μ*g/mL, respectively (Fig. 6, all model predictions of the mini-ensemble measurements can be found in the SM). These values are consistent with previous estimates of *n* = 10-13 myosin heads (from ∼20-25 myosin heads/*μ*m (68) at 25*μ*g/mL times ∼\0.5*μ*m of actin available in the laser trap (69)).

In the *in vitro* motility measurements, we also do not know the exact number of myosin molecules present. We therefore performed a series of simulations with different ensemble sizes (see Methods). In these simulations, we simulate myosin molecules interacting with an actin filament and record the position of the actin filament over time, *x* (*t*). We fit a straight line to *x*(*t*) and the slope is the actin speed. An ensemble of *n* = 40 gave good agreement between our motility measurements and the best-fit parameter set for two of the fibers (fibers 15 and 17), yet the other parameter set (fiber 18) predicts nearly two-fold higher speed at saturating ATP (see SM). However, simulations with myosin ensembles composed of a mixture of myosin, each of which has parameters from one of the three best-fit parameter sets with equal probability, gives reasonable agreement with our measurements (Fig. 6), consistent with the observation that slower myosin molecules have a disproportional effect on *in vitro* motility speed (70).

Overall, we found the agreement between our molecular measurements and the model predictions to be good. This result suggests not only that the model reasonably captures the molecular interactions between myosin and actin, but also that our measurements are self-consistent, i.e. that the interactions between myosin and actin are sufficiently similar in all of the measurements to be described by a single mechanism.

### Force redevelopment following ramp shortening

Despite the model’s success in capturing force during ramp shortening, force during and after step lengthening/shortening, the history dependence of isometric force, and molecular measurements, the model consistently over-predicts the force rise after the shortening ramp (Fig. 5A), even as it captures the force rise after the shortening step in the step lengthening/shortening experiments (Fig. 5B). One possibility for the discrepancy between model and measurement is sarcomere non-uniformity. Previous muscle models have shown emergent behavior in multi-sarcomere simulations (13, 71, 72), so it seems plausible that immediately after the shortening ramp, sarcomeres in the fiber might adjust their lengths and produce different forces than the model, which assumes that all sarcomeres in the fiber act identically. To test this idea, we performed Monte-Carlo simulations of multiple sarcomeres (five half sarcomeres) in series undergoing a ramp shortening of 5% over 0.1s and found that they predict a force response consistent with 1) a single half sarcomere with the same number of myosin, 2) five half sarcomeres in parallel, and 3) our differential equation model. We find that, in agreement with previous modeling work (73), thick filament activation stabilizes half sarcomeres and results in uniform shortening (Fig. 7, inset). We therefore conclude that, in the model, sarcomere non-uniformity is unlikely to contribute significantly to muscle force for the experiments considered here.

**Figure 7.**
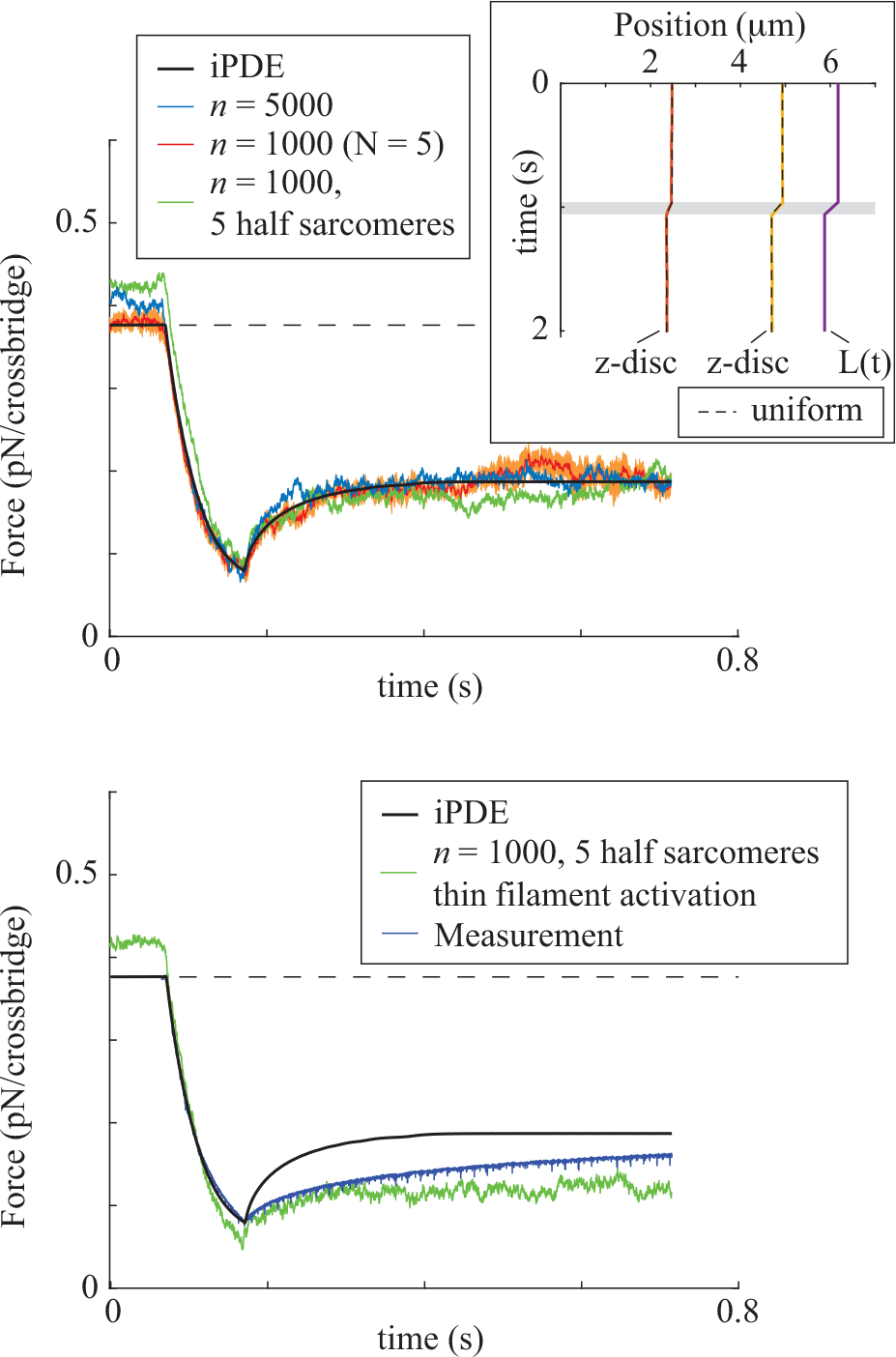
Comparison of the model iPDEs to Monte-Carlo simulations. (Top) iPDE model agrees with Monte-Carlo simulations of multiple sarcomeres in parallel and in series. Inset shows a kymograph of sarcomere length in the simulations with 5 half sarcomeres in series, solid colored lines, compared to uniform sarcomeres, dashed lines. (Bottom) Differences appear between the iPDE and Monte-Carlo simulations when thin filament activation is added. Like in our measurements, the rate of force redevelopment after shortening is reduced with thin filament activation. Parameters from fiber 17 used in all simulations.

A second possibility for the discrepancy between model and measurement is thin filament activation. In particular, during many of the ramp shortening experiments, force drops to < 20% of isometric. If there is an associated drop in the number of attached cross-bridges, myosin’s attachment to actin would then be decreased because of a lack of strongly bound myosin to keep the thin filament fully active via strong-binding (74). This would occur even at high calcium, because myosin’s attachment to actin is reduced roughly 2-fold in the presence of regulatory proteins (19). Simulations of five half sarcomeres with thin filament activation, implemented as in Longyear et al. 2017 (19), result in a different force response than predicted by the iPDE model and, suggestively, a slower rate of force redevelopment after the shortening ramp (Fig. 7). We therefore conclude that capturing the slower force rise after shortening in our model will likely require both thin and thick filament activation mechanisms.

## CONCLUSIONS

Multiscale models of muscle function require data sets from the same tissue across multiple scales so that model parameters may be estimated and model predictions tested. Molecular and cellular measurements are technically challenging, so that few labs are experts in both and measurements are generally performed under conditions that optimize the success of experiments at one scale or the other. Here, we present a molecular-to-cellular data set, all measured with muscle harvested from the same source (rabbit psoas) and all performed under similar conditions. To demonstrate the utility of these data, we developed a (to our knowledge) new muscle model, based on the PDE formalism of Huxley (28) that includes thick filament activation whereby myosin enter (or leave) a folded, self-inhibited state if force on the thick filament drops below (or climbs above) a threshold. We used parameter optimization to fit the model to a subset of our cellular measurements. The model then reasonably predicts the remaining cellular measurements (Fig. 5). Remarkably, the model also reasonably predicts the molecular measurements (Fig. 6). Though several studies have used fits to molecular measurements to reasonably predict fiber measurements, e.g. (19, 75), we are not aware of another study where molecular measurements have been quantitatively predicted from model fits to fiber measurements. Finally, because the model fits both the fiber’s force response to shortening and stretch, which are thought to have history dependencies of differing molecular origin (76), the model suggests a unifying molecular mechanism of muscle’s history dependence. The possibility of our model both quantitatively describing muscle measurements from the scale of single molecules to whole cells and also explaining muscle’s history dependence, which has lacked a widely-accepted explanation for 70 years, warrants further exploration.

## MATERIALS AND METHODS

To ensure a self-consistent data set, we used psoas muscle from New Zealand White Rabbits, housed at the University of Calgary. Single cell measurements were performed on skinned fibers at the University of Calgary. For the *in vitro*, molecular scale measurements, muscle from the same animals was shipped to the University of Massachusetts, where myosin was then isolated and the experiments performed. The two labs coordinated to match buffers and other experimental conditions as closely as possible. Here, we provide details of these experiments and details of our modeling. Additional, more technical details and model validation may be found in the Supplementary Material.

### Single fiber experiments

#### Sample preparation and experimental setup

Single skinned fibers isolated from psoas muscle of six-month-old female New Zealand White Rabbits were used in this experiment. Rabbits were euthanized by an intravenous pentobarbital (240mg/mL) injection. Strips of psoas muscle were harvested and tied to wooden sticks before storing in rigor solution (see Solutions) at 4°C. Muscle strips were transferred into a rigor overnight and glycerol (50:50) solution within 4 to 5 hours. On the next day, muscle strips were moved to a rigor and glycerol (50:50) solution and kept at −20° C for 10 to 14 days for skinning (77, 78). Ethics approval for all procedures was granted from the University of Calgary Animal Ethics Committees.

On the experimental day, a small piece of psoas muscle was cut from the skinned muscle strips and transferred to a petri dish containing relaxing solution with protease inhibitors. Single fibers were isolated from the muscle piece and one end of the fiber was tied to a suture before transferring it to the testing chamber. After the fiber was moved to the testing chamber, one end of the fiber was attached to a length controller (322C, Aurora Scientific Inc., ON, Canada) and the other end to a force transducer (403A, Aurora Scientific Inc., ON, Canada) to allow measuring the lengths and forces during the experiment. Fibers were set at an average sarcomere length of 2.4*μ*m using a He-Ne laser. All experiments were performed at 15° C.

### Force-velocity experiments

Fibers (*n* = 8) were maximally activated at an average sarcomere length of 2.4*μ*m and shortened by 5% of the initial fiber length (to an average sarcomere length of 2.28*μ*m) at various average sarcomere speeds (12*μ*m/s, 4.8*μ*m/s, 2.4*μ*m/s, 1.2*μ*m/s, 0.48*μ*m/s, 0.24*μ*m/s, and 0.12*μ*m/s) in a random order (Fig. 1A). Some fibers, *n* = 3, were shortened at very slow speed (0.004*μ*m/s) and others *n* = 2, where shortened at a faster speed (24*μ*m/s).

### Quick stretch tests

Fibers (*n* = 8) were maximally activated at an average sarcomere length of 2.4*μ*m using an activating solution (pCa =− log_10_ ([Ca^2+^]) = 4.2). After force reached the steady-state, fibres underwent quick stretches of 0.5% and 0.75% of the initial fibre length in 0.2 ms followed by shortening to the original length (Fig. 1B).

### Solutions

Rigor solution: Tris 50mM, potassium chloride 100mM, magnesium chloride 2mM, ethylene glycol-bis (2-aminoethylether)-N,N,N’,N’-tetraacetic acid (EGTA) 1mM at pH 7.0.

Rigor overnight solution: Tris 50mM, potassium chloride 2mM, magnesium chloride 2mM, sodium chloride 100mM, and EGTA 1mM at pH 7.0.

Relaxing solution: Potassium propionate (Kprop) 170mM, 3-(N-morpholino) propanesulfonic acid (MOPS) 20mM, magnesium acetate (MgAc_2_) 2.5mM, K_2_EGTA 5mM, adenosine triphosphate (ATP) 2.5 mM at pH 7.2.

Activating solution: Kprop 170mM, MOPS 10mM, MgAc_2_ 2.5mM, CaEGTA 5mM, ATP 2.5mM at pH 7.2 (78).

### In vitro experiments

#### Proteins

Myosin used in all the experiments was purified from rabbit psoas muscle as previously described (79), with minor modifications. Actin was purified from chicken pectoralis muscle as previously described (80). Actin was also fluorescently labeled with 100% tetramethylrhodamine (TRITC)/phalloidin for in vitro motility and 50% TRITC/phalloidin and 50% biotin/phalloidin for the laser trap assay experiments. Regulatory proteins, purified human cardiac tropomyosin and rabbit fast skeletal troponin (complex of TnI/TnC/TnT), were obtained from Life Diagnostics (West Chester, PA) and maintained in a −80° C freezer until the day of the experiment.

### Solutions

All buffers were brought to the appropriate pH using a calibrated pH-meter (Fisher Scientific). pH was adjusted as needed with 1 M potassium hydroxide (KOH) to increase the pH or decreased with 1 M Acetic Acid (CH_3_COOH). All individual buffer ion concentrations, total ionic strength of 95mM, and free Ca^++^ were determined using WinMaxC (81). Prior to each motility experiment, the isolated myosin was mixed with unlabled actin and 1 mM ATP in a high-salt buffer and then ultracentrifuged at 400,000g for 20 minutes at 4° C to remove myosin unable to hydrolyze ATP (i.e., dead head spin down). In regulated thin filament motility, the free Ca^++^ concentration was set to pCa 4 using appropriate amounts of CaCl_2_ added to the motility buffer, with the solution recipe determined using the stability constants and equations within the WinMaxC software program (81). The ATP concentration was varied from 10*μ*M to 2 mM for the motility experiments.

The single molecule experimental buffer was as described above, and additionally had 1*μ*m streptavidin-coated silica microspheres (Bangs Labs Inc.), and three separate [ATP] (1, 2.5, and 5 *μ*M). The mini ensemble experimental buffer also had the silica microspheres and 100*μ*M ATP.

### In vitro motility assay

Flow cells were used to allow each element to be sequentially injected into the chamber, which was maintained at 30° C as in previously established methods (82). The solutions were loaded sequentially as follows: 35 *μ*L of isolated monomeric rabbit skeletal myosin (0.100 mg/mL) in a high salt myosin buffer (300 mM NaProprionate, 25 mM MOPS, 1 mM EGTA, 4 mM MgAcetate, pH 7.4, 1 mM DTT) was incubated for 30 seconds. Following this, 35 *μ*L of bovine serum albumin (BSA) (0.5 mg/mL) in actin buffer (25 mM NaProprionate, 25 mM MOPS, 1 mM EGTA, 4 mM MgAcetate, pH 7.4, 1 mM DTT) was loaded and incubated for 60 seconds. 35 *μ*L of unlabeled actin fragments in the absence of ATP was loaded and incubated for 30 seconds. 35 *μ*L of ATP wash (1 mM ATP in actin buffer) was loaded and incubated for 30 seconds. 35 *μ*L of TRITC-labeled actin (10nM in actin buffer) was loaded and incubated for 30 seconds, and repeated for a total of 70 *μ*L. This was followed by 35 *μ*L of actin buffer. Finally, 35 *μ*L of motility buffer (a 1:1 ratio of 1% methylcellulose and actin buffer with 95mM total ionic strength with the appropriate addition of NaProprionate, the appropriate amount of ATP for the given experiment, and an oxygen scavenging system) was loaded into the flow cell. The total ionic strength was calculated using the Debye-Huckel equation as previously described (47).

Regulated in vitro motility was performed similarly. However, following loading the 70 *μ*L of fluorescently labeled actin filaments, 35 *μ*L of reconstitution solution containing actin buffer with 0.25 *μ*M tropomyosin and 0.75 *μ*M troponin were added and allowed to incubate in the flow cell for 7 minutes to reconstitute the regulated thin filaments (82, 83). In addition, an excess of 100 nM tropomyosin and 100 nM troponin was added to the final motility buffer to ensure full regulation of the thin filaments was maintained (84).

To visualize the fluorescently labeled actin filaments, the flow cells were then placed onto a Nikon Ti-U (Nikon, Tokyo, Japan) inverted microscope with a 100X, 1.4 numerical aperture CFI Plan Apo oil-coupled objective and an ICCD camera (Stanford Photonics, Inc., Palo Alto, CA). Videos of separate fields were captured, for each flow cell, at 10 frames୶s^−1^ for each condition. Images were captured by a Epix-LVDS frame grabber (Epix, Inc., Buffalo Grove, IL) which was coupled to an ICCD camera using Piper Control™ 2.5 software (Stanford Photonics, Inc. Palo Alto, CA).

#### Analysis of in vitro motility data

Actin filament velocities and the percentage of moving filaments were determined using an automated filament tracking algorithm within the imageJ plugin WRMTRK (85). Filaments with a velocity lower than 0.13*μ*m/s were considered stationary and filaments shorter than 0.5*μ*m were eliminated from analysis. The mean of all the actin filament velocities in the video was taken as the average velocity and then each of the videos were averaged for the individual flow cell. For each condition, typically 3-5 flow cells were used to generate the average velocity data with at least 3 videos per flow cell.

#### Laser trap assay

The three-bead laser trap assay was used to determine the duration of strong binding to an actin filament and the displacement generated by single molecules and mini ensembles of our rabbit psoas myosin. Myosin was adhered to a nitrocellulose-coated microscope slide where 3 *μ*m silica microspheres served as pedestals for myosin to attach. The myosin concentration was adjusted to yield either single molecule (0.2 *μ*g/mL) or mini ensemble (25 and 50 *μ*g /mL) binding events. Following the addition of myosin, the flow cells were incubated for 5 minutes with bovine serum albumin (0.5 mg/mL) to inhibit non-specific interactions between the actin filament and the nitrocellulose-coated coverslip surface.

A three-axis piezo-controlled stage (Mad City Labs, Inc.) was maneuvered to attach a biotin/TRITC-labeled actin filament to two 1*μ*m neutravidin-coated silica microspheres (Bangs Labs, Inc.) held in two time-shared optical traps. Once an actin filament was attached to the 1*μ*m microspheres, the filament was extended between the optically trapped microspheres, to a apply pretension of 3-4pN to the actin filament. The bead-actin-bead dumbbell assembly was lowered into close proximity of the 3*μ*m myosin-coated pedestal bead. The displacements generated by actomyosin binding activity were tracked from the output from quadrant photodiode via interferometry at a sampling rate of 5 kHz, as previously described (22).

#### Single molecule data processing and analysis

Displacement records from the single molecule laser trap assay were analyzed using custom programs within R (v4.04, R Core Team). A two-state Hidden-Markov Model (86) was used for binding event identification using running mean and variance transformations of the raw displacement data from the laser trap assay, and changepoint analysis was used to determine the start and end of each binding event as previously described (22).

#### Mini-ensemble data processing and analysis

The displacement records form the mini-ensemble laser trap assay were analyzed using custom programs within R (v4.04, R Core Team) to automatically detect events, determine peak displacements, as well as event duration, as previously described (19, 47). Briefly, actomyosin binding events were determined after baseline correction by applying a low pass filter using a running mean with a 10ms window and a displacement > 8nm above baseline applied to the running mean used to mark the start and end of events. Once detected, the event duration was determined as the total time a displacement magnitude of >8nm above baseline, with time-off the duration between the end of an one event and the beginning of the next.

### Fitting fiber data with the model

In order to fit the model to our measurements of force in the skinned fiber experiments, we 1. numerically solved the iPDEs (Eqn. 1) subject to the two constraints of force balance and conservation of probability, 2. simulated two ramp shortening experiments for a given parameter set, 3. simulated a step stretch/step shortening experiment for a given parameter set, 4. compared simulation results to measurements, 5. performed parameter optimization to maximize goodness-of-fit. We now describe each of these steps. All code was written in Matlab (Mathworks, Natick, MA) and is available upon request.

#### Numerical solution to iPDEs

We solve the following iPDEs to simulate the ramp shortening and step stretch/shortening experiments:

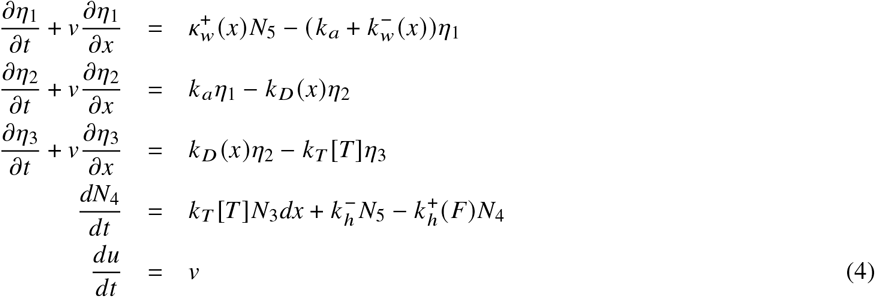

Note the slight difference from Eq. 1 in that we do not solve the equation for *N*_5_ (but rather calculate it from conservation of probability as described below) and have an additional equation to keep track of the extension of the contractile element,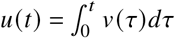.

Our general approach to solving the iPDEs is to first turn them into a system of integro Ordinary Differential Equations (iODEs) using the method of characteristics. This is accomplished by defining the characteristic
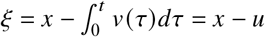. Then, each partial differential

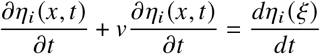

There are an infinite number of these equations, one for each value of *x*. We therefore discretize *x* into *x*_1_, *x*_2_, *x*_3_, …, and solve a system of iODEs at each *x*_*i*_.

To solve the iODEs, we use a fourth-order Runga-Kutta method with a constant step size. We use this method as opposed to variable step size methods implemented in Matlab (e.g., ode45) because at each time step we must integrate each *η*_*i*_ to find *N*_*i*_, calculate force *F*, and perform a root find to determine the sliding velocity, *v*, that balances forces to satisfy Eq. 2. To find *N*_*i*_ we numerically integrate *η*_*i*_, for *i* = 1, 2, 3, using the trapezoid method (as implemented in Matlab’s trapz function). *N*_4_ comes directly from the solutions of Eq. 4, and we find *N*_5_ from conservation of probability, *N*_5_ = 1− (*N*_1_+*N*_2_+*N*_3_+*N*_4_). To find *F* and to find *v* with the root find, we must account for the fact that myosin molecules transition between the folded, inhibited state and the unfolded, active state.

Each probability density, *η*_*i*_ (*x, y, t*), is a function of three variables, *x* the molecular extension, *t* time, and *y* a variable that defines the force at which myosin transitions from the folded, inhibited state to the unfolded, active state. Similarly, each probability, *N*_*i*_ (*y, t*) is a function of two variables. However, there is no explicit *y* dependence in the equations and the equations at different *y* are coupled only through the force *F*. We can therefore discretize *y* into *y*_1_, *y*_2_, *y*_3_, … and, at each time step, perform a Runga-Kutta step as described above for every *y* _*j*_, with force *F* (*t*), sliding velocity *v* (*t*) and binding probabilities *N*_*i*_ (*t, y* _*j*_) calculated from the previous time step. Then, to calculate force, we numerically calculate

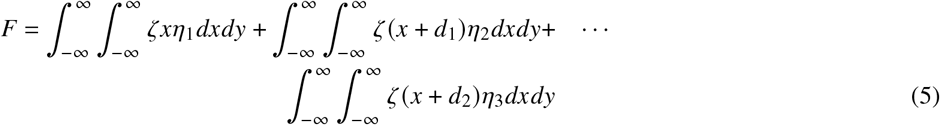

by first using the trapezoid method to integrate in *x*, and then using the right-endpoint method to integrate in *y*. Using this force calculation, we can perform a root find to determine *v*(*t*) to satisfy Eq. 2, which depends on the length of the fiber, *L*(*t*). We accomplish this root find using Matlab’s fsolve function.

Simulations start from isometric steady-state, which we calculate by solving Eq. 1 with *v* = 0 and time derivatives set to 0(see Supplement). Then, the basic steps to numerically simulate the iPDEs are 1. perform a root find to determine *v* (*t*+Δ*t*), given *L*(*t*), *η*_*i*_ (*x, y, t*) and *N*_*i*_ (*y, t*), 2. calculate *η*_*i*_ (*x, y, t*+Δ*t*) (*i* = 1, 2, 3), *N*_4_ (*y, t*+Δ*t*), and *u*(*t*+Δ*t*) using the value of *v* (*t*+ Δ*t*), 3. perform numerical integration to determine *F*(*t*+Δ*t*) and *N*_*i*(_ *t*+ Δ*t*) (*i* = 1, 2, 3, 5).

In all simulations, we used constant time and space steps (Δ*t*, Δ*x*, and Δ*y*). We fixed Δ*y* = 0.01, though we used slightly different Δ*t* and Δ*x* for the ramp shortening and step stretch/shortening experiments (see below). To increase the robustness of our simulations, we implemented a cutoff so that the force-dependent reaction rate 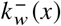 had a maximum value of 100000s^−1^. This cutoff allows us to take a larger time step in the simulations and decreases numerical error by keeping the equations non-stiff. Additionally, rather than implementing the folding/unfolding transition as a step function (Eq. 3), we use a smooth function (the error function). We similarly smooth out the step in fiber velocity in the ramp shortening experiments and the step in fiber length in the step stretch/shortening experiments (see below). Finally, in an effort to decrease numerical error in our simulations we set the tolerances for the root find to 1 · 10^−10^.

These simulations take fiber length as a function of time, *L* (*t*), as an input and give force as a function of time as an output. Additionally, the simulations require the specification of the seven model parameters (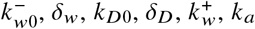). Simulating 0.15s of two ramp shortening experiments and 0.2s of one step stretch/shortening experiment takes about 45 minutes on a personal computer. For comparison, simulating an ensemble of 5000 myosin molecules in one ramp shortening experiment using Monte Carlo methods (see description, below) takes several hours on the same computer.

#### Ramp shortening

For the ramp shortening experiments, we discretize *x* into 1000 equally spaced points between −13nm and 13nm (Δ*x* = 0.026). We used a time step of Δ*t* = 2 ·10^−5^s. The velocity step was smoothed using an error function, which we then integrated to get *L* (*t*).

#### Step stretch/shortening

For the step stretch/shortening experiments, we discretize *x* into 500 equally spaced points between −9nm and 9nm (Δ*x* = 0.036). We used a time step of Δ*t* = 2.5 · 10^−5^s. Length steps were smoothed using an error function.

#### Comparing model to data

The model systematically over-predicts the force rise after the ramp shortening experiments (Fig. 5A). This error likely arises from an effect that is not yet included in the model, e.g. thin filament activation (see Discussion). Fits that minimize the sum of the squared difference between model and measurement compensate for the over-predicted rise in force by systematically under-predicting force during shortening. In short, the predicted force-time plot is bad everywhere, rather than just after shortening.

We avoided this problem by fitting the time derivative of force. This procedure has the additional advantage that the rate of force development/loss is related to actomyosin kinetics. As many of the rate constants governing these kinetics are parameters of the model, fitting the time derivative of force will result in more robust estimates than fitting force itself.

Another problem that arises from under-predicting force during shortening relates to the force-velocity relationship and actin filament speed in the motility assay. In particular, unloaded shortening velocity measured from force-velocity relationships in muscles and muscle fibers is typically faster than actin filament speed in the motility assay (87). A viable multiscale model should capture this effect. However, a model that under-predicts force in the force-velocity relationship will naturally predict a slower unloaded shortening velocity. Force-velocity curves can be constructed by measuring the steady-state force achieved during a shortening ramp (e.g. (64)) and, while our measurements do not achieve steady-state, the force at the end of the ramp shortening shows similar qualitative behavior and can be used to construct a force-velocity curve (see SM). We therefore wanted the model to faithfully capture the force at the end of the ramp shortening, so we added the squared difference between model and measurement at this point to the cost function.

The cost function was therefore the sum of the mean squared difference between modeled and measured force rates for 0.15s of two ramp shortenings (duration 25 and 100ms). The measured force rate was determined by first smoothing the force measurement with a moving window of width 10 points and then numerically calculating the derivative with a forward difference. In addition, the cost function included 10^4^ times the difference between modeled and measured force at the end of the shortening ramp. Finally, the cost function included 10^4^ times the mean squared difference between modeled and measured force for 0.2s of the step lengthening/shortening experiments. The factor of 10^4^ accounts for the fact that typical forces (relative to isometric) are of order 10^0^, while typical force rates are of order 10^2^, and the cost function measures the squared difference.

### Optimization of fit

To minimize the cost function, we used a combination of Bayesian optimization and a Nelder-Mead simplex method. Each optimization started with an initial random seed for the seven parameters, with initial guesses selected from a uniform distribution in the following ranges 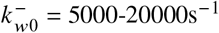, δ_*w*_ = 0.04-16nm, *k*_*D*0_ = 1000-7000s^−1^, δ_*D*_ = 0.01 2nm, 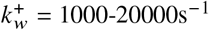, *k* _*a*_ = 10-800s^−1^, and ζ_*s*_/*F*_*iso*_ = 0.1-1nm^−1^ where *F*_*iso*_ is the isometric force per myosin. We then ran 30 iterations of Bayesian optimization, using Matlab’s Bayesopt function to identify an initial best-fit parameter set. We then used these initial best-fit parameters as a seed for an optimization using the Nelder-Mead simplex method, Matlab’s fminsearch function. We ran this for ∼ 130 iterations. Overall, optimization took around a week of simulation time (see SM for an example optimization and a justification for terminating the optimization before convergence).

### Predicting in vitro data with the model

We simulated single molecule, mini-ensemble, in vitro motility, and fiber measurements using a modified Gillespie algorithm (14, 19). Briefly, the simulations keep track of the state and molecular extension of each myosin molecule, and the times of each possible chemical reaction are picked randomly from an appropriate probability distribution. The minimum of these times is selected, the state of the given molecule updated, and extension of each myosin molecule and position of actin are updated by assuming that the system is in mechanical equilibrium. In simulations with the laser trap, the lasers apply a force on the actin filament equal to the trap stiffness times the displacement. In the motility assay, force on the actin filament is zero. Except for the simulations with multiple sarcomeres (described below) in the fiber simulations the displacement of actin is defined so we omit this step and simply update the extension of all attached myosin molecules each time step.

#### Single molecule

For the single molecule simulations, each time step we add random noise from the Brownian motion of the bead-actin-bead dumbbell (normally distributed, with mean 0 and standard deviation 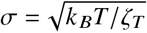, where ζ_*T*_ = 0.04pN/nm is the stiffness of the laser trap). We additionally add a small amount of normally distributed random noise (mean 0, σ = 2nm) to model noise in the laser trap. Simulated data were then saved and analyzed using the same custom code we used to analyze the single molecule measurements (see above).

#### Mini ensemble

Mini ensemble simulations were the same as the single molecule, except that we add cooperativity that occurs because the binding of the first myosin molecule restricts the z-fluctuations of the bead-actin-bead dumbbell. We account for this by having the first myosin molecule attach at 0.01 times the subsequent values (i.e., the weak binding attachment rate for the first molecule is 0.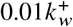). Simulated data were analyzed using the same algorithm we used to analyze the mini ensemble measurements (see above).

#### In vitro motility

The output of the motility simulations is actin position as a function of time. We ran the simulations for 100000 time steps, or around 0.5s depending on parameters. To estimate actin velocity, we fit a straight line to the position as a function of time, interpolated to have equally spaced time points, and the velocity was the slope of this line.

#### Solution ATPase

When in solution, myosin experiences no external force. Each rate constant that is force-dependent in the model therefore takes it’s force-independent value (e.g., *k*_*D*_(*x*) is replaced with *k*_*D*0_, 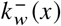 is replaced with 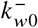, and so on). Additionally, the iPDEs (eq. 1) become ODEs. Finally, since we are interested in steady-state ATPase, all of the time derivatives in eq. 1 are zero, and we have a homogeneous system of linear equations. To solve these equations, we calculate the nullspace of the matrix defining the linear system (we do this numerically in Matlab) and then normalize to ensure conservation of probability, which allows us to solve for each *N*_*i*_ (*i* = 1, 2, 3, 4, 5). Then, ATPase 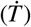 is the flux through the cycle, which we calculate by 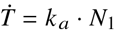.

### Monte-Carlo fiber simulations

In the iPDE simulations, we assume that each half sarcomere starts with the contractile element in isometric steady-state and each parallel spring with zero extension. However, if titin binds to actin in a muscle fiber, presumably there is some connection between titin binding and myosin binding to actin (77). To attempt to simulate this, we started our Monte-Carlo simulations from 0 myosin bound with the stiffness of the parallel spring (ζ _*p*_) set to 0. Because the simulations took a long time to run (e.g., a 1s simulation took about 24 hours), we applied a small stretch to increase the number of attached myosin and then engaged the parallel spring. For the multi-sarcomere simulations, this introduces non-uniformity, since each parallel spring has a different rest length. We ran these simulations for ∼6s prior to starting the ramp stretch to ensure that the system was in steady-state. This procedure results in some variability in the initial isometric force, which is why there is an initial difference in force between the Monte-Carlo and iPDE simulation in Fig. 7.

## AUTHOR CONTRIBUTIONS

SW designed the research, with assistance from WH and ED. VJ performed preliminary experiments to help develop the experimental protocol for the muscle fiber experiments. SL performed the cellular experiments. CM and ED performed the laser trap experiments. MW performed the in vitro motility experiments. SW developed the mathematical model and performed all simulations. BS developed and performed the data analysis of the laser trap measurements. TL assisted in cellular experiments and in harvesting muscle. SW wrote the article with feedback from all authors.

## ACKNOWLEDGMENTS

This work was supported, in part, by the Canada Research Chair program and the Nigg Chair for Biomechanics Research to SY and VJ and by NIH R01-GM135923. The authors are grateful for helpful discussions with Manoj Srinivasan on optimization and numerics, and with Mike Previs on thick filament activation.

## SUPPLEMENTARY MATERIAL

An online supplement to this article can be found by visiting BJ Online at http://www.biophysj.org.

## Notes

### Competing Interest Statement

The authors have declared no competing interest.

