## Supplemental Material for "Modeling Thick Filament Activation Suggests a Molecular Basis for Force Depression"

S. Liu, et al.

This supplement contains additional details of our measurements, data selection and analysis, and modeling.

#### 1 Skinned fiber measurements

##### 1.1 Initial isometric force

In our experiments, isometric force showed history dependence. Each ramp shortening and step stretch was performed from a single activation, so it is possible that the results of our measurements depend on the order in which they were performed. As one way of minimizing this effect on our results, we randomized the order in which the shortening/stretches were performed. In addition, the fiber was given five minutes to recover in between length perturbations so that isometric force did not change appreciably over the course of the experiment (Fig. S1)

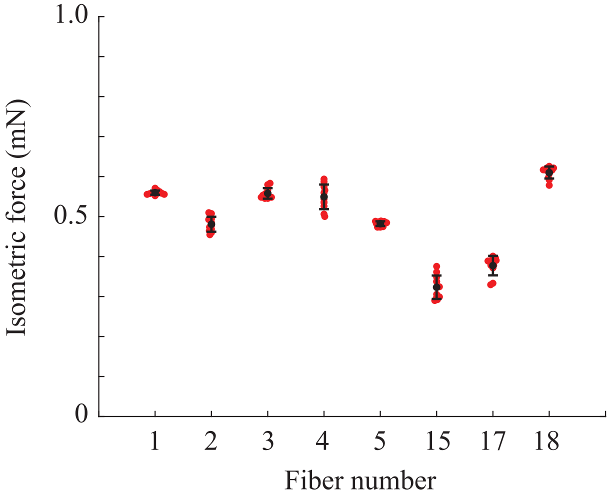


Figure S1: Isometric force recovers between length perturbations. Force 200ms prior to each step stretch or ramp shortening, averaged over 500 points (25ms) is plotted for each fiber. Red dots are individual measurements in a beeswarm plot where lateral displacement indicates density; black dots are mean and error bars show standard deviation. Variation in isometric force is less than 10% with a mean value of 4%. Fibers 15, 17, and 18 were fit by the model.

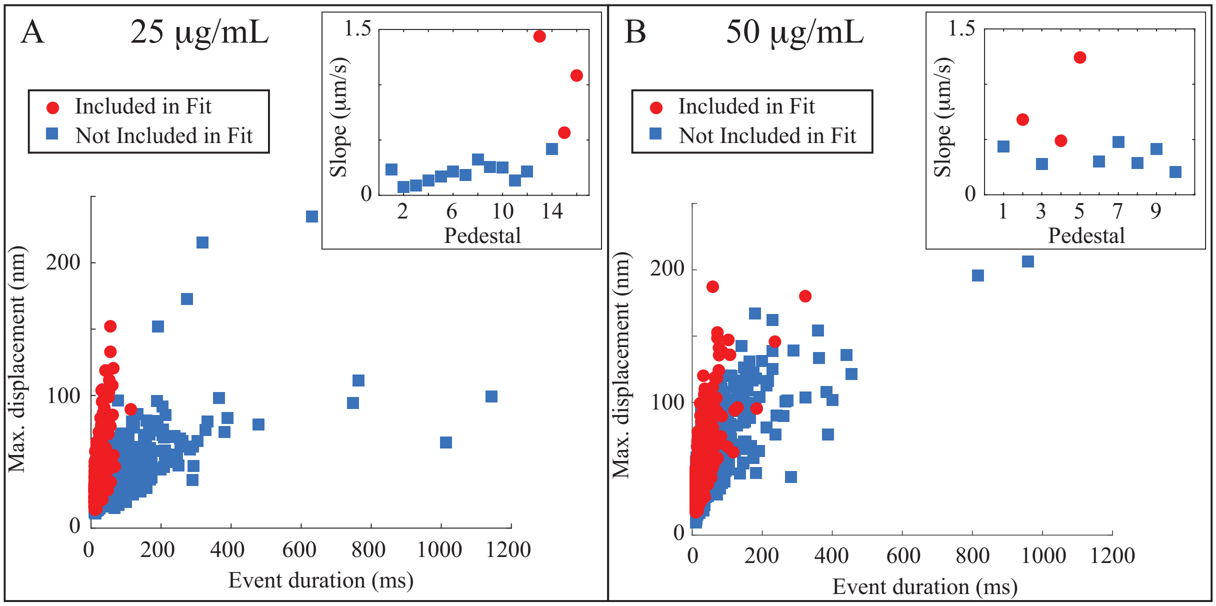


Figure S2: Eliminating potentially confounding effects from the mini ensemble data. By fitting a plot of maximum event displacement vs. event duration with a straight line for each pedestal (inset shows slope), we removed data from pedestals that produced long-lasting, low displacement events. A. shows the lower myosin density (25 $\mu$g/mL) and B. shows the higher myosin density (50 $\mu$g/mL).

### 2 Molecular experiments

#### 2.1 Selection criteria for mini ensemble measurements

We performed more mini ensemble laser trap measurements than are presented in the main text. A subset of these measurements showed long-lasting binding events with minimal displacement. When included in our analysis, these events give a decrease in average event lifetime as myosin concentration is increased from 25 to 50$\mu$g/mL (Table S1). It is difficult to understand how an increase in the number of myosin interacting with actin in the mini ensemble assay would decrease event lifetime.

Table S1: Mini-ensemble measurements

| Myosin concentration ($\mu$g/mL) | $d_{max}$ (nm) | $t_{event}$ (ms) |
|---|---|---|
| 25 ($n = 2242$) | $32.58 \pm 0.34$ | $35.99 \pm 1.21$ |
| 25 ($n = 617$) | $38.22 \pm 0.70$ | $20.08 \pm 0.45$ |
| 50 ($n = 3000$) | $42.32 \pm 0.39$ | $30.16 \pm 0.75$ |
| 50 ($n = 813$) | $47.12 \pm 0.76$ | $23.41 \pm 0.73$ |

Numbers listed are mean plus/minus SEM.

There are a variety of ways that event lifetimes can be prolonged, including interactions between actin and the surface of the pedestal and/or the presence of dead-heads or otherwise damaged myosin. Such effects are not included in the model. To minimize these potentially confounding effects, we plotted event lifetime and maximum displacement for our measurements at each pedestal (interactions at a pedestal resulted in $\sim 200$ binding events). We then fit these with a straight line and selected only data from the three pedestals with the highest slope, which corresponded to a slope above $\sim 500$nm/s (Fig. S2). This subset of the data showed increased displacement and decreased attachment time compared to all of the data, and also showed increases in both displacement and attachment time as myosin density was increased

(Table S1).

### 3 Fitting the model to data

To optimize the fit of the model to our measurements, we used a combination of Bayesian optimization (Matlab's bayesopt function), followed by a simplex-based optimization (Matlab's fminsearch function, see Methods). We ran the simplex-based optimization for $\sim 130$ iterations (around a week of simulation time), stopping the optimization prior to convergence. We stopped the optimization because, after $\sim 100$ iterations, there was little change in the goodness of fit (Fig. S3B).

In the analysis presented in the main text, we assume that these partial optimizations give similar results as a full optimization in 1) predicting the results of molecular-scale experiments, and 2) predicting the fiber measurements not included in the cost function, i.e., that Figs. 5 and 6 of the main text would remain approximately the same if we allowed the optimization to fully converge. Moreover, we argue that the model is predictive, but we do not quantify its predictive ability. Finally, some of the best-fit parameters presented in Table 2 of the main text vary from fiber to fiber. E.g., the unloaded ADP release rate, $k_D^0$, ranges from 3130 to 7300s$^{-1}$. This variability suggests that either the fits are insensitive to some parameters or that each fiber requires a different set of parameters. We performed a series of additional optimizations in order to justify our assumptions, quantify the model's predictive ability and show that the model fits are insensitive to some parameters.

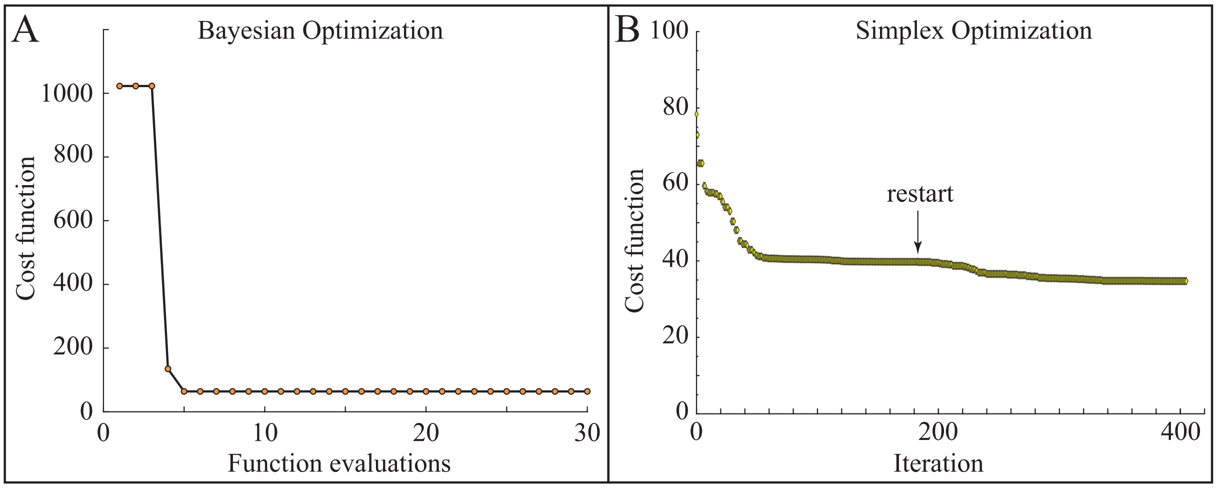


Figure S3: Optimization trajectory for fits to measurements from Fiber 18. A. Bayesian optimization. B. Simplex optimization. Note the different scales in A and B. In B, the arrow indicates where the simplex optimization converged prematurely, and we restarted the optimization.

#### 3.1 Partial vs. full optimization

To evaluate how terminating the optimization affected our results, we ran an optimization (for Fiber 18) to convergence. We obtained convergence after more than 400 iterations (Fig. S3). The best-fit parameters obtained from the full optimization were generally different from those obtained from the partial optimization (see Table S2).

#### 3.2 Parameter sensitivity

Because of the computational expense in getting the optimization to converge, we chose one parameter to investigate in a sensitivity analysis. We were particularly interested in how the model results depend on the

Table S2: Best fit parameters for full and partial optimization to measurements from Fiber 18

| | Full Optimization | Partial Optimization |
|---|---|---|
| $k_{D0}$ ($s^{-1}$) | 8713 | 7300 |
| $\delta_D$ (nm) | 1.52 | 1.61 |
| $k^-_{w0}$ ($s^{-1}$) | 100000* | 12600 |
| $\delta_w$ (nm) | –* | 3.54 |
| $k^+_w$ ($s^{-1}$) | 2541 | 12600 |
| $k_a$ ($s^{-1}$) | 529 | 296 |
| $\zeta_s$ (pN/nm/myosin) | 0.0230 | 0.0208 |

*Optimal $k^-_{w0}$ exceeded cutoff of 100000$s^{-1}$ required for numerical stability (see Methods).

unloaded ADP release rate ($k^0_D$), since that was highly variable in our fits to different fiber measurements (Table 2 of the main text), and because it is thought to limit sliding velocity in the motility assay (e.g. Walcott et al. 2012). We therefore did a sensitivity analysis for $k^0_D$.

To do so, we took advantage of the fact that the optimization converged prematurely after ∼180 iterations (Fig. S3B, arrow). At that point, $k^0_D$ was 2535$s^{-1}$ while it was 8713$s^{-1}$ at the end of the optimization. We therefore re-ran optimizations to convergence from this point, but with $k^0_D$ fixed at $1000, 2535, 4000$, and 6000$s^{-1}$. For all but the slowest ($k^0_D = 1000s^{-1}$), which did not fit the data as well, all values of $k^0_D$ gave similar best-fits (i.e., within 15%) when compared to the optimal (Fig. S4A). We therefore conclude that the fits are insensitive to $k^0_D$ if it is above a critical value $\approx 2000s^{-1}$.

##### 3.3 Molecular predictions

In our analysis, we assumed that full and partial optimizations will give similar predictions of molecular-scale experiments. This assumption is true provided that parameters that produce near-optimal fits to the data will all produce similar molecular-scale predictions. While we can test this simply by comparing the fully and partially optimized fits, the sensitivity analysis provided us with a way to more fully test this assumption. In particular, we re-ran simulations of the mini-ensemble and in vitro motility experiments (Fig. 6B and C of the main text) with each best-fit parameter set from our sensitivity analysis, with $k^0_D$ fixed at $1000, 2535, 4000$, and 6000$s^{-1}$ and also the best-fit parameters with $k^0_D = 8713s^{-1}$. From these, we estimated 1) the number of independent myosin heads present in the mini ensemble (Fig. S4B) and 2) in vitro motility speed as a function of ATP (Fig. S4C).

If our assumption is correct, then the values of $k^0_D$ that produce near-optimal fits to the data (2535, 4000, 6000, and 8713$s^{-1}$ ) will all produce similar molecular-scale predictions and these will be similar to the predictions of the partial optimization. Indeed, this is what we found, with all predicting $N$ = 11-14 independent myosin heads in the mini ensemble experiments for the lower myosin density (25$\mu$g/mL) and $N$ = 15-17 independent myosin heads for the higher myosin density (50$\mu$g/mL), Fig. S4B. Similarly, they all predicted similar motility speed vs. ATP plots, Fig. S4B. We also observed that the parameter set that did not fit the data well ($k^0_D = 1000s^{-1}$) made different predictions for the molecular experiments, Fig. S4B, C. This result shows that the molecular predictions are not a generic property of the model, but rather suggests that the molecular predictions are a generic property of models that successfully fit the cellular measurements. We therefore found support for our assumption that the partial optimizations give similar results as a full optimization in predicting our molecular experiments.

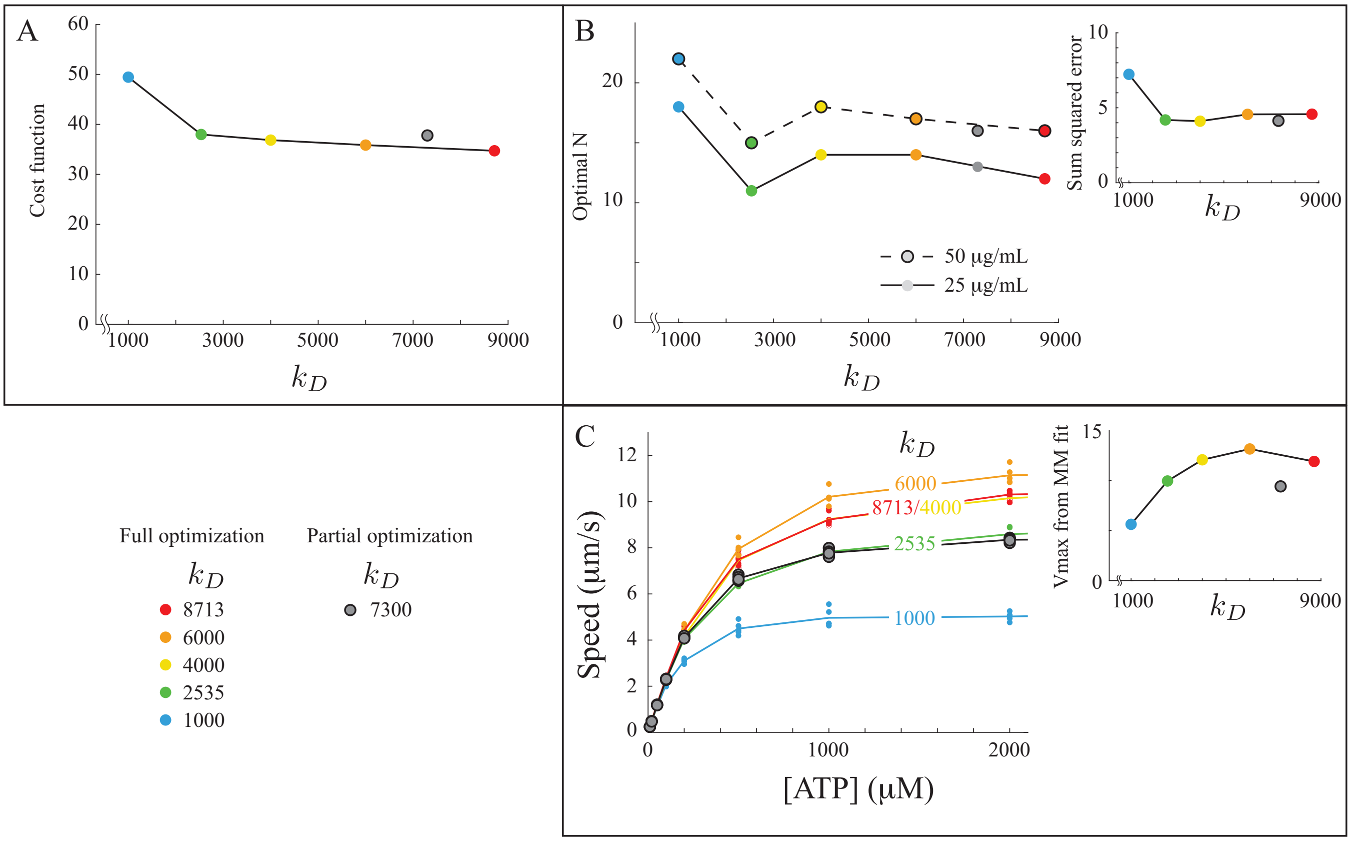


Figure S4: Molecular predictions of the model are similar for good fits to the cellular measurements. A. A sensitivity analysis showing the cost function of the best-fit of the model to measurements from fiber F18 for different values of the unloaded ADP release rate, $k_D^0$. Over a broad range ($2535 < k_D^0 < 8713\mathrm{s}^{-1}$), the model generates similar fits to the measurements, whereas the fit becomes worse for $k_D^0 = 1000\mathrm{s}^{-1}$. B. Good fits to the cellular measurements predict similar numbers of myosin heads in the ensemble, while the worst fit ($k_D^0 = 1000\mathrm{s}^{-1}$) predicts more myosin heads. Good fits to the cellular measurements fit the mini ensemble data better than the worst fit (inset). The predicted number of heads from the partial optimization (from the main text) is similar to the predicted number for the good fits. C. Good fits to the cellular measurements predict similar velocities in the motility assay, while the worst fit ($k_D^0 = 1000\mathrm{s}^{-1}$) is different. This result suggests ADP release does not necessarily limit sliding velocity in the motility assay in the model. Inset shows maximum sliding velocity as predicted by a Michaelis-Menten fit to the simulated motility data and also shows that the good fits to the cellular data predict similar results at the molecular scale.

##### 3.4 Cellular predictions

We assume that full and partial optimizations will give similar predictions of cellular scale experiments. That is, we only fit a subset of the cellular measurements performed on a given fiber and we assume that the predictions of the remaining fiber measurements for the partial optimization will be similar to those for the full optimization. Moreover, although the predictions we observed from the partial optimization (Fig. 5 of the main text) appear good, we did not quantify the goodness-of-fit. Here, we use our full optimization and the optimizations from the sensitivity analysis to support this assumption and to quantify the model predictions.

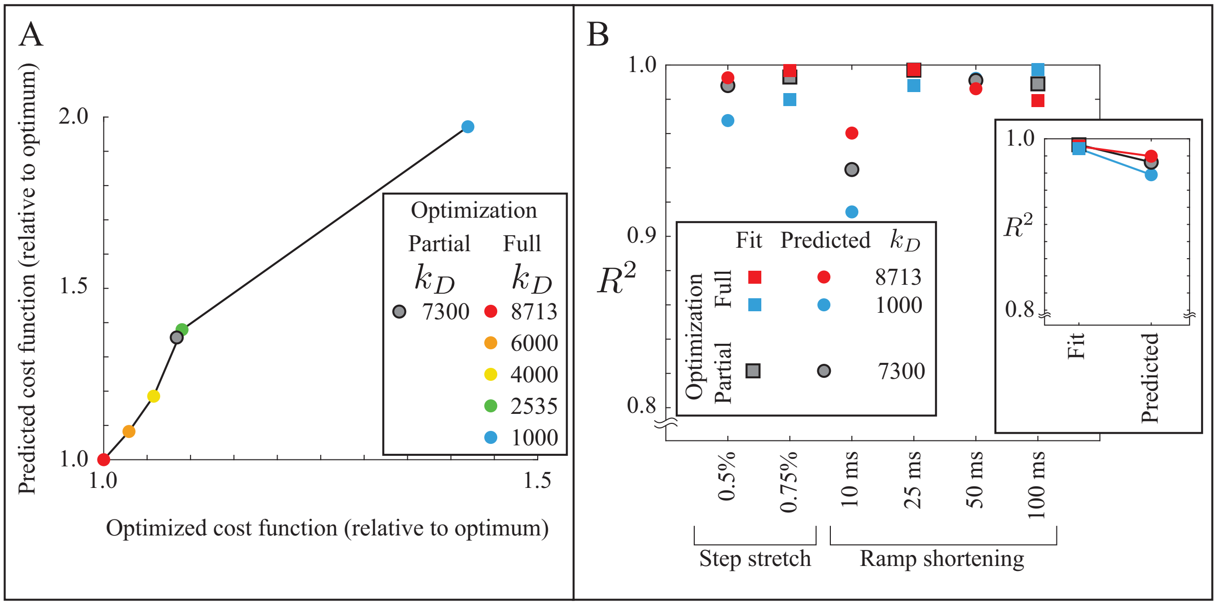


Figure S5: Model predictions of cellular measurements improve as optimization-of-fit converges. A. Cost function of three measurements not included in the optimization (step stretch of 0.5%, ramp shortenings of 10ms and 50ms) vs. the cost function of the three measurements in the optimization (step stretch of 0.75%, ramp shortenings of 25ms and 100ms). A positive slope indicates a positive correlation, i.e. that an increase in the cost function in the optimization results in an increase in the cost function for the measurements not included in the optimization. B. Assessing the goodness-of-fit to the force measurements shows that as the fit to the measurements improves (squares) the predictions of the model get more accurate (circles). Inset shows the average of all three measurements. Note that the $R^2$ calculation for the shortening ramp includes the force until the end of shortening, while the step stretch includes 0.2s of data (as shown in Figs. S6-S9).

If the model is predictive, then we would expect that as the optimization-of-fit converges, the model will more successfully predict the remaining measurements. That is, if the cost function for each fiber measurement is minimized by the same parameters, then the cost function for any measurement is optimized as the fit for one measurement is optimized. To test this idea, we plotted the cost function for the three measurements for which the fit was optimized (step stretch of amplitude 0.75%, ramp shortening over 25 and 100ms) vs. the cost function of three measurements which were not included in the optimization (step stretch of amplitude 0.50%, ramp shortening over 10 and 50ms). We observe a strong correlation. As the goodness-of-fit of the optimized cost function decreases $\sim$ 40%, the goodness-of-fit of the non-optimized measurements decreases $\sim$ 100% (Fig. S5A).

To quantify goodness-of-fit, we calculated the $R^2$ value of the fits to the step stretch experiments and the fits to the ramp shortening experiments prior to the isometric hold after shortening. We excluded the fits after the isometric hold because the model does not fit force redevelopment after shortening. We find that, as the optimization converges, the $R^2$ of both the optimized and predicted measurements increase (Fig. S5B). Moreover, though improvement in the fit for the optimized measurements is modest, the improvement for the predicted measurements is larger (Fig. S5B, inset). For the fully optimized fits, the predictions have an average $R^2 = 0.98$, indicating that the predictions are successful. Finally, for the partially optimized fits, the predictions have an average $R^2 = 0.97$, suggesting that the partial optimization predicts the measurements nearly as well as the full optimization.

##### 3.5 Conclusion

This detailed analysis of fits to measurements for Fiber 18 shows that, at least for that measurement, the partial optimization gives similar results as the full optimization in predicting molecular-scale experiments (Fig. S4) and predicting the fiber measurements not included in the cost function (Fig. S5). We additionally show that the fits are not sensitive to some parameters (Fig. S4A), explaining why there is variability in the best-fit parameters in Table 2 of the main text. Together these results suggest that the optimum in parameter space is large and shallow, so that the optimization rapidly finds good fits but then only slowly converges to the best fit (as shown in Fig. S3B). Using a partial instead of a full optimization therefore allows us to rapidly find good parameters without having to devote the computational expense to discover the optimal parameters, and these good parameters give similar predictions both at the molecular and cellular scale as the optimal parameters.

#### 4 Model assumptions

##### 4.1 Extrapolation of $f(y)$

To estimate $f(y)$, which defines how many myosin are active as a function of steady-state force, we used our measurements from very slow (30s) 5% shortening ramps (a shortening speed of 0.00167 FL/s). In so doing, we do not need to assume a functional form of $f(y)$ and eliminate the need of any parameters to define it. However, this procedure only defines $f(y)$ for forces achieved during the slow shortening ramps, where force drops to a minimum of between 50-60% isometric (Fig. 4C of the main text). To predict $f(y)$ for the smaller forces that occur during the faster shortening ramps, we therefore had to extrapolate. We chose to use an error function to extrapolate, since it gave about 20-30% myosin active in the absence of force (e.g., [1]). In preliminary simulations, we tried a few different choices of extrapolation functions, including linear and quadratic. For some simulations, the system was unable to sustain activation, and this appeared to happen less frequently if $f(y)$ was convex (as in the error function extrapolation). We therefore chose to use the error function extrapolation in all of our simulations.

##### 4.2 Near-maximal activation at isometric force

In preliminary simulations, we found better agreement between model and the fiber measurements when the myosin was not completely activated under isometric conditions. In the simulations, 93% of the myosin is active under isometric conditions and full activation only occurs when force rises above isometric, as during the step stretch experiments. This assumption is therefore important for the step stretch experiments, but not for the shortening ramp experiments where force is always below the initial isometric force. To quantify the importance of this effect, we ran an optimization (to convergence) with full activation for isometric conditions and found that the optimal value of the cost function was higher (46.04) than the cost function with 93% of the myosin active at isometric (34.72). Therefore, some additional activation during lengthening improves the fit of the model to the data.

### 5 Model results

#### 5.1 Fits and predictions from all three fibers

In Figure 5 of the main text, we compare the model to measurements only of Fiber 17. In this section, we show the remaining model fits (Fiber 15 Fig. S6, Fiber 17 Fig. S7 and Fiber 18 Fig. S8 and Fig. S9). In these figures, we show the data that were fit by the model (i.e., force rate in the ramp shortening experiments) as well as the minimum force achieved during the shortening ramp, which is related to the steady-state force-velocity relationship but is not identical because our measurements do not reach steady state. For Fiber 18, we include both the partial optimization as well as the full optimization.

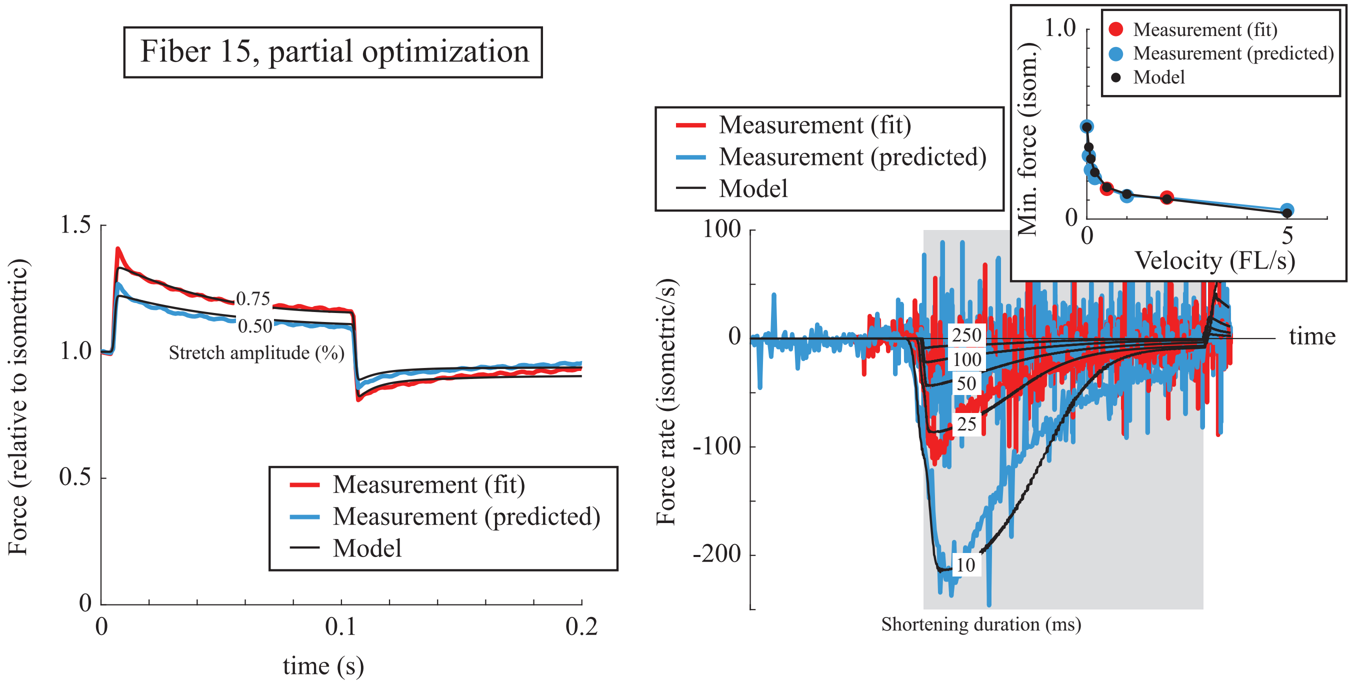


Figure S6: Cellular fits and predictions for fiber F15. In all plots, measurements fit by the model are in red, while measurements predicted by the model are in blue. The force measured during step lengthening/shortening is shown on the left (cf. Fig. 1B of the main text). The force rate (the time derivative of force) is shown for ramp shortening on the right. Note that time is rescaled so that the shortening ramp is initiated and ended at the same point, indicated by the gray box, for each of five measurements (shortening durations of 10, 25, 50, 100, and 250 ms). Inset shows the minimum force at the end of the ramp shortening for different shortening rates (converted to shortening velocities). The cost function combines the mean squared error of the force-time trace in the step shortening/lengthening, the mean squared error of the force-rate time trace in the ramp shortening, and the minimum force at the end of the ramp shortening, as described in the Methods section of the main text.

#### 5.2 Molecular-scale predictions from all three fibers

In Figure 6 of the main text, we compare the model to molecular measurements using only the best-fit parameters of Fiber 17 for the mini-ensemble and using a 1:1:1 mixture of molecules with parameters from Fibers 15, 17, and 18 for the in vitro motility assay. Moreover, in the mini-ensemble measurements, we show histograms of event lifetime and maximum displacement for both the model to our measurements, but the fits we used to predict the number of independent myosin heads were calculated from cumulative probability plots. Here, we show the model and measurements for both the mini-ensemble and in vitro motility assays for all three best-fit parameter sets (those for Fibers 15, 17, and 18). In addition, we show the cumulative probability plots for the mini-ensemble (Fig. S10).

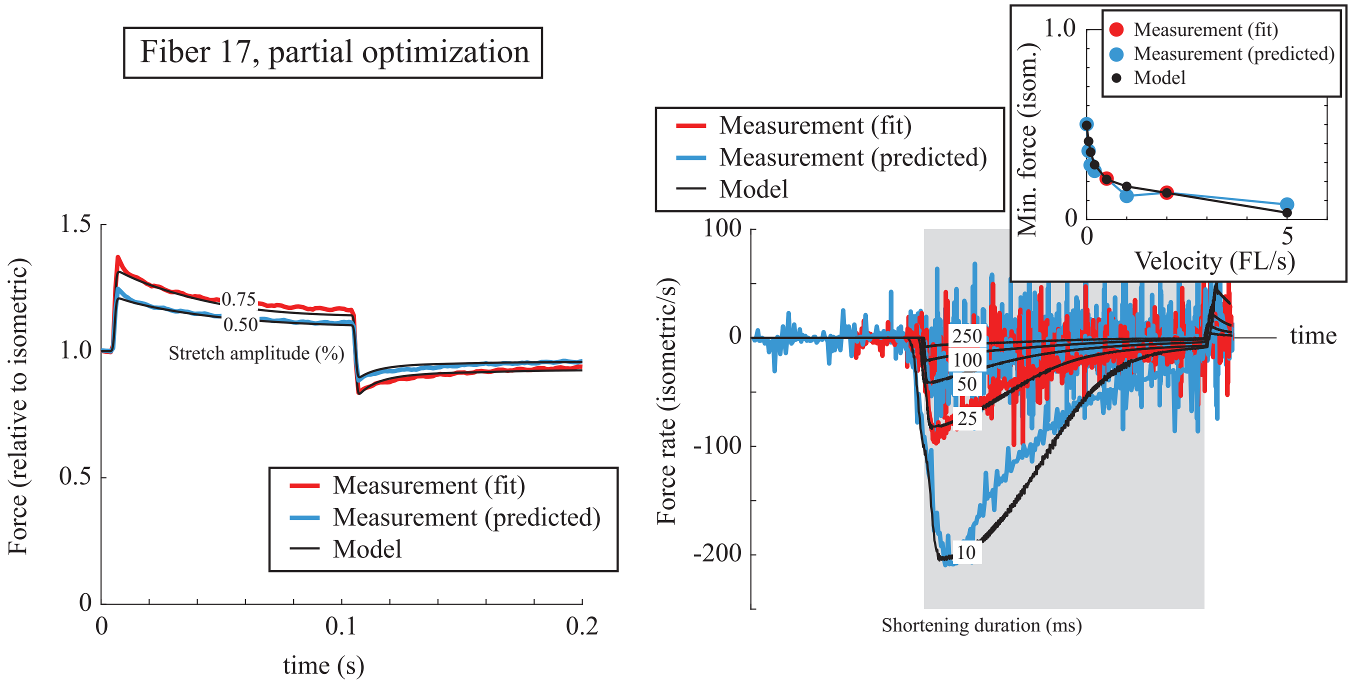


Figure S7: Cellular fits and predictions for fiber F17. See Fig. S6 for a description.

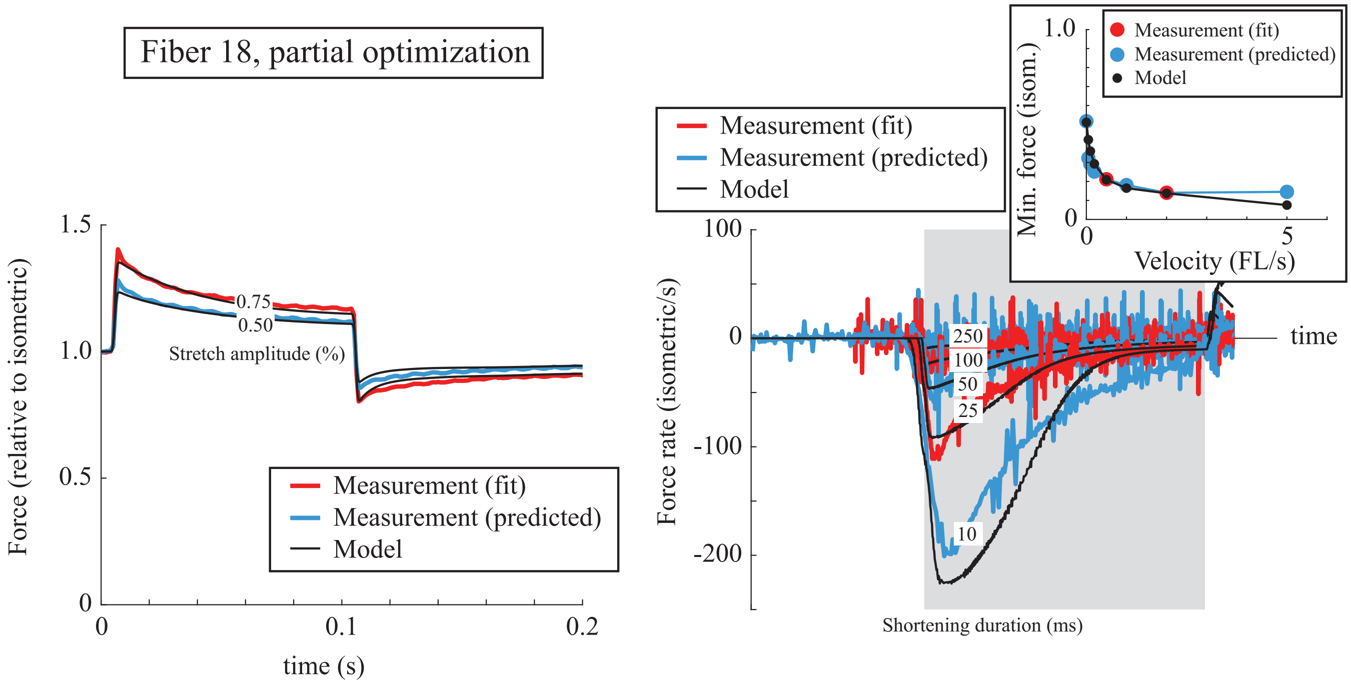


Figure S8: Cellular fits and predictions for fiber F18. See Fig. S6 for a description.

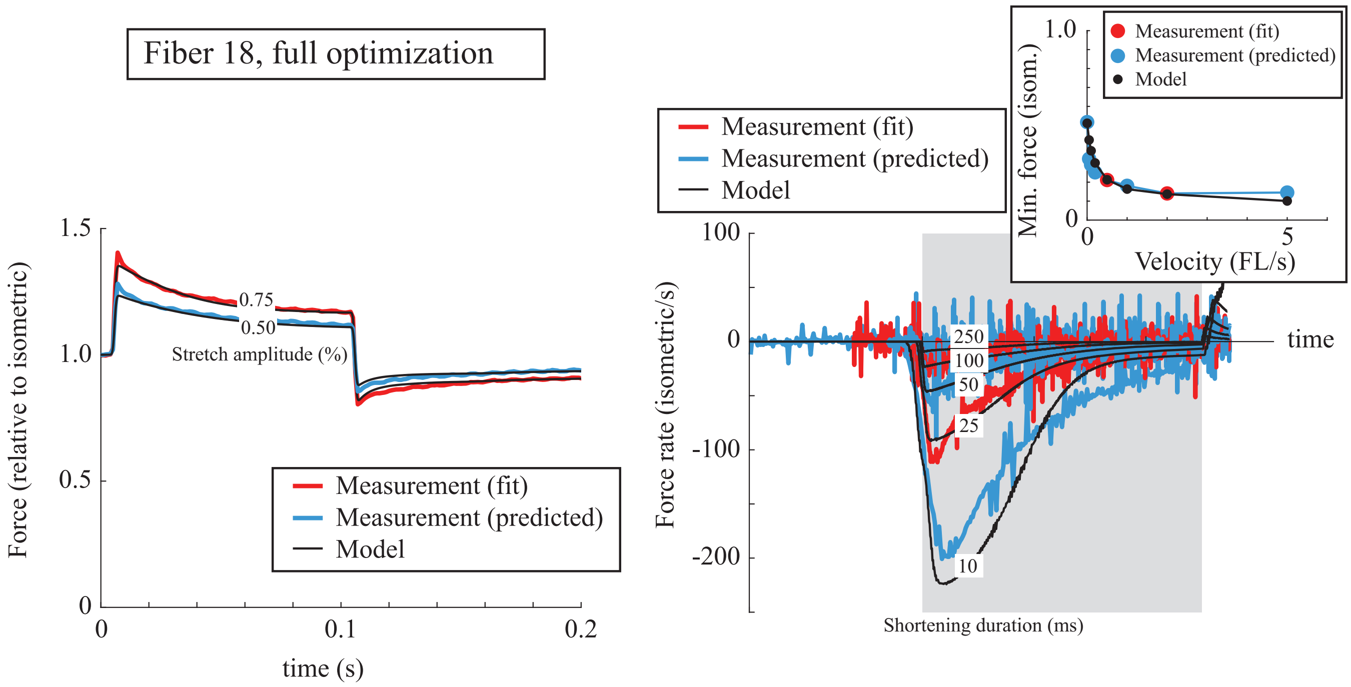


Figure S9: Cellular fits and predictions for fiber F18, where the optimization converged. See Fig. S6 for a description.

The predictions of in vitro motility (Fig. S10A) are variable, with maximum values between 5 and 10$\mu$m/s. The experimental data are more consistent with the slower speeds. The mini ensemble measurements are more consistent, with all parameter sets predicting nearly identical numbers of myosin at each of the two myosin densities, and nearly identical distributions of maximum displacement and lifetime of the binding events (Fig. S10B).

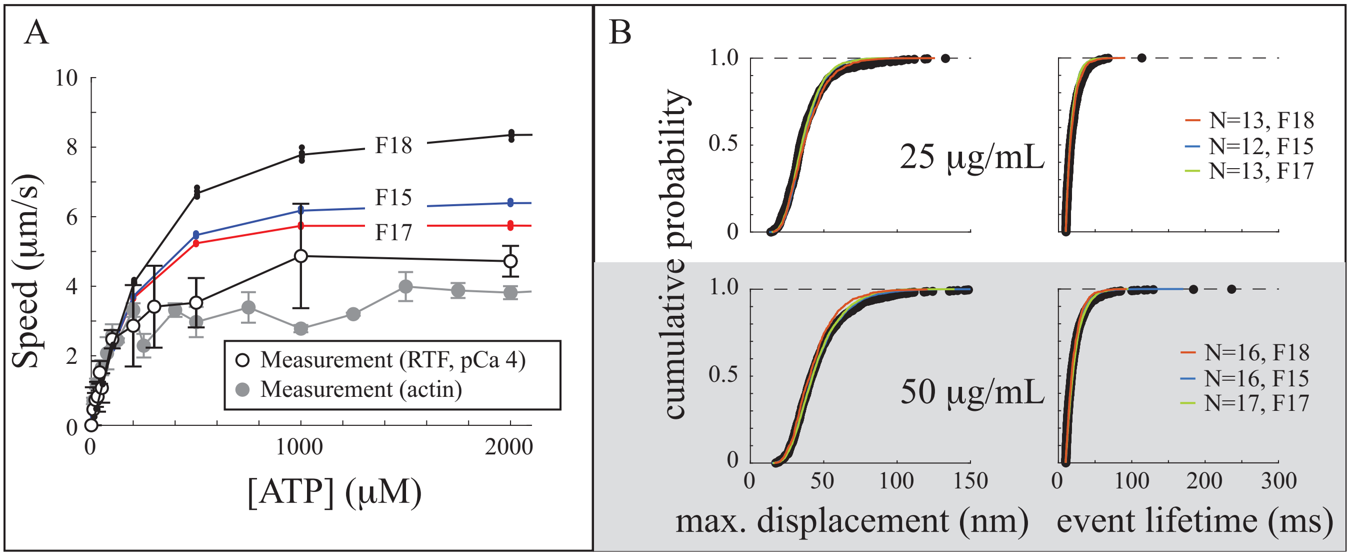


Figure S10: Predictions of molecular measurements for all three fits. A. In vitro motility as a function of ATP concentration for regulated thin filaments (hollow dots) at high calcium and for actin (gray dots) are similar to, but a bit slower than the predictions from the three fits. The predicted curves are variable, with maximum predicted sliding velocities between 6 and 8 $\mu$m/s. Error bars show standard deviations. B. Maximum displacement and event lifetime for mini-ensemble measurements (black dots) are similar to model predictions. The predicted curves are all similar and predict similar numbers of independent myosin heads in the experiments. Lower myosin density (25$\mu$g/mL) is shown in the top plots, and higher myosin density (50$\mu$g/mL) is shown in the bottom plots.
